# Quantifying competition between two demersal fish species from spatiotemporal stomach content data

**DOI:** 10.1101/2024.04.22.590538

**Authors:** Max Lindmark, Federico Maioli, Sean C. Anderson, Mayya Gogina, Valerio Bartolino, Mattias Sköld, Mikael Ohlsson, Anna Eklöf, Michele Casini

## Abstract

Inference on competition is often made on indirect patterns of potential competition, such as population trends and spatiotemporal overlap in diet and distribution. However, these indicators do not test if the contested resources are limited in supply, nor if they decline as competitor biomass increases. Using stomach content and biomass data, we evaluate food competition between Atlantic cod (*Gadus morhua*) and flounder (*Platichthys* spp.) in the Baltic Sea. We quantify diet overlap and fit geostatistical mixed models to evaluate effects of local-scale covariates on stomach contents. The dietary overlap is low and does not decline with predator density. We find that cod feed less on the isopod *Saduria entomon* at high flounder densities. However, the total prey weight in cod is not affected by flounder densities. This suggests interspecific food competition is not limiting the overall feeding of cod, but affects its diet composition. In addition, we find support for intraspecific food competition in large cod and flounder. Our study illustrates the importance of local-scale processes when inferring competition from stomach content data.

## Introduction

Competition occurs when individuals — whether of the same or different species — interact for access to the same limited resource that improves their fitness; the concept has been a long-standing central tenant of ecological theory (Volterra 1926, Smith 1935, Elton 1949, Macarthur and Levins 1967, Schoener 1983, Holomuzki *et al*. 2010). Competition affects population dynamics via individual-level processes such as growth, dispersal and survival (Andersen 2020). On a community level, competition regulates species composition via competitive exclusion and partitioning of resources in space or time (Chesson 2000), and on an evolutionary level it drives trait diversity (Grant and Grant 2006). Competition has mostly been studied using mathematical models (Macarthur and Levins 1967, Lotka 1978, Tilman 1982) or by experimentally manipulating the abundance of a potential competitor in either a laboratory or field setting (Paine 1966, Werner and Hall 1977, Larson 1980, Goldberg and Barton 1992, Schoener 1983, Link and Auster 2013).

In natural systems, much of the empirical work on competition has been done on relatively small spatiotemporal scales, such as intertidal zones, freshwater lakes, or forest patches and on more or less sessile organisms using manipulative approaches (Paine 1966, Werner and Hall 1977, Lubchenco 1978, Link and Auster 2013). However, numerous organisms, including many marine fishes, are widely distributed and undergo seasonal migrations, as well as ontogenetic habitat and diet shifts (Heincke 1913, Stevenson *et al*. 2022, Ciannelli *et al*. 2022), such that the relevant spatiotemporal scales are much larger than what is covered by much of the seminal experimental work on competition. In such cases, competition is often inferred using statistical models fitted to observational spatial or temporal data of abundance or biomass (Jennings and Kaiser 1998, Link 2002, Ovaskainen *et al*. 2017, Godefroid *et al*. 2019). Despite the limitations of such correlative approaches, numerous studies have identified strong effects of interspecific interactions in marine food webs, where the majority of food web links are often weak compared to terrestrial ecosystems (see reviews in Jennings and Kaiser 1998, Minto and Worm 2012). Studies have often leveraged the fact that commercial fishing alters species composition and relative abundance within marine fish communities, thus constituting a form of semi-controlled manipulation experiment (Jennings and Kaiser 1998, Szuwalski *et al*. 2017), either via removal of predators, which can increase competition among prey, or by removal of potential competitions, by targeting specific species. However, it is difficult to confidently attribute patterns in population abundance to competition (Schoener 1983, Goldberg and Barton 1992, Jennings and Kaiser 1998, Dormann *et al*. 2018), since population dynamics of marine fishes are heavily influenced by stationary and non-stationary environmental drivers acting on different time scales, as well as by other biotic interactions such as predation, prey-dependent survival, and indirect interactions (Jennings and Kaiser 1998, Ohlberger *et al*. 2022).

A more direct test of competition is by relating some measure of fitness, such as growth, or body condition to densities of potential competitors. An early example of this is the strong decline in growth in adult North Sea plaice following the Second World War when the stock biomass increased as a result of the reduced fishing mortality (Margetts and Holt 1948, Rijnsdorp and van Leeuwen 1992). As for interspecific competition, Casini *et al*. (2006) found that the decline in body condition of sprat and herring were largely driven by the increase in biomass of the sprat stock in the Baltic Sea. In the Baltic Sea, potential competition has also been identified between Atlantic cod (*Gadus morhua*) and flounder (*Platichthys flesus* and *Platichthys solemdali*), in that they exhibit opposite population trajectories and high spatiotemporal overlap (Orio *et al*. 2019). The latter has also increased over time, as there has been a spatial contraction of the distribution of cod and an increase in the spatiotemporal overlap with flounder (Orio *et al*. 2019). However, Lindmark *et al*. (2023), did not find effects of cod or flounder density on the body condition of cod, and flounder density in the southern Baltic was higher in the late 1980s when cod condition was still high (Orio *et al*. 2017, Lindmark *et al*. 2023). This suggests intra- and interspecific competition is not particularly strong, assuming body condition is related to food availability and by extension the abundance of potential competitors.

Analysis of stomach contents can prove useful to more directly look for signals of competition compared to traits such as condition and growth, which are shaped by complex interactions between environmental and fishing effects, and are also built up over longer time periods. Combining diet data with a body growth model, Neuenfeldt *et al*. (2020) showed that food limitation of cod between 21–35 cm is at an all time high in the Baltic Sea. The latter could be a result of dwindling benthic prey availability (Orio *et al*. 2019, 2020, Neuenfeldt *et al*. 2020), due to spatial mismatch between cod and benthic prey (Lindmark *et al*. 2025), or competition with flounder who feed predominantly on benthic prey (Haase *et al*. 2020), although the latter remains to be tested. Moreover, food competition can be manifested in multiple ways beyond reduced feeding rates on shared prey. It can also alter prey choice—if preferred prey decline under high competition, predators may switch to less profitable alternatives. These shifts may leave dietary fingerprints that can be tracked with metrics, such as Schoener’s overlap (Schoener 1968), measuring diet similarity between groups (Barnes *et al*. 2018). Detecting signals of competition in total feeding rates, diet diversity, and overlap metrics can thereby help to expose the mechanisms that redistribute energy in food webs and determine who wins or adjusts when resources become increasingly limited.

In this study, our aim is to advance our understanding of intra- and interspecific food competition, using Baltic cod and flounder as a case study. The productivity of eastern Baltic cod is at a historical low due to a reduced body growth (Mion *et al*. 2021), body condition (Casini *et al*. 2016b, Lindmark *et al*. 2023) and increased natural mortality (Casini *et al*. 2016a). Hence, it is critically important to further understand factors affecting cod fitness. Identifying competitive interactions in non-manipulative systems requires a diversity of approaches, using multiple indicators at different spatiotemporal scales (Pianka 1976, Link and Auster 2013). Therefore, we apply a suite of models at different spatiotemporal scales. First, we quantify diet overlap between cod and flounder overlap (a prerequisite for food competition) by exploring basin-wide dietary similarity using model-based ordination. Next, we estimate to what degree the biomass density of the potential competitors affects diet overlap, as would be expected under the hypothesis that competition drives diet separation. Lastly, we move beyond dietary overlap and more directly test which factors (abiotic and biotic) explain variation in the weights of prey in predator stomachs. This is done by fitting spatiotemporal generalized linear mixed models (GLMMs) to local-scale stomach content data (weight of prey in individual predator stomachs) in relation to the local density of potential competitors and abiotic covariates.

## Methods

### Data

Cod and flounder stomachs were collected by the Swedish University of Agricultural Sciences, Department of Aquatic Resources, Sweden, during the bi-annual Baltic International Trawl Survey (BITS) conducted in the first and fourth quarter of the year following the BITS protocol (ICES 2017). Flounder refers to Baltic flounder (*Platichthys solemdali*), and European flounder (*Platichthys flesus*), which are morphologically indistinguishable (Momigliano *et al*. 2018). We used data from the southwestern Baltic Proper (Fig. 1) between 2015 and 2022 (data from quarter one in 2019 were not available), because during this period both cod and flounder stomachs were collected. The stomach sampling was designed to collect flounder and cod stomachs from the same trawl hauls. Whenever possible, one flounder and one cod stomach were collected for each 1 cm of fish length and trawl haul (see Fig. S1 for size distributions by year). Stomachs were extracted and frozen as quickly as possible by staff onboard. Frozen stomachs were sent to the National Marine Fisheries Research Institute in Gdynia, Poland, for taxonomic identification of prey organisms to the lowest level possible and for calculation of total prey weights. The number, size and weight of individual prey were also measured when possible. Regurgitated stomachs were removed during data processing. After data processing, 5982 cod stomachs (3209 in the first quarter and 2773 in the fourth quarter), and 3882 flounder stomachs (2093 in the first quarter and 1789 in the fourth quarter) were available for analysis.

**Figure 1:**
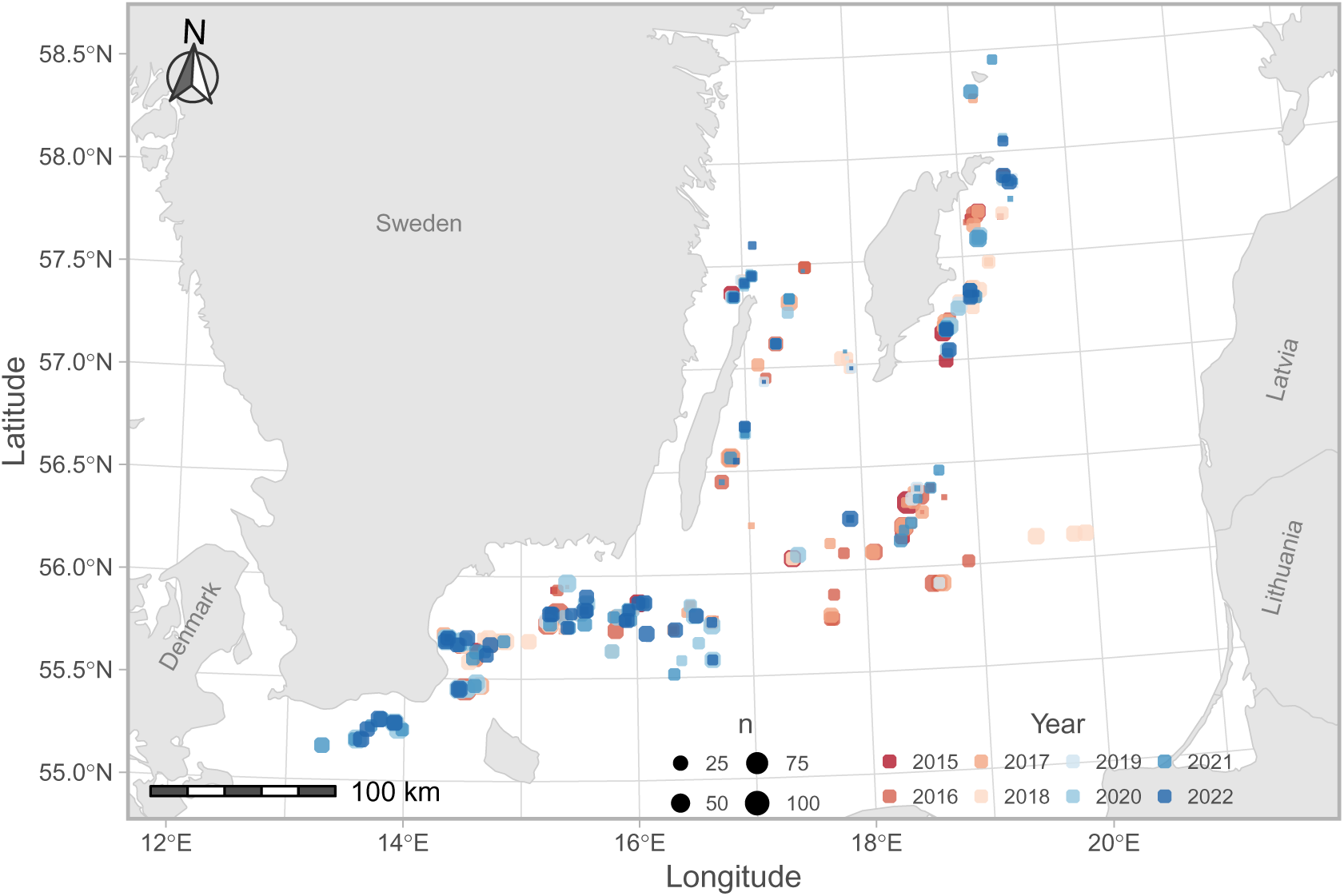
Location of stomach samples in the southern Baltic Sea. Colors correspond to years and the size (area) of the point corresponds to sample size by year at that location.

Because stomach samples stem from length-stratified sampling, samples from hauls with large catches represent a larger fraction of the population than samples from hauls with small catches. To account for this potential source of bias, we included sampling (or “design”) weights either in our models or when calculating mean diets if that was the response variable. The weights were constructed for each observation by calculating the ratio of the number of fish caught of the predator group to the number of fish sampled in the associated haul, and then scaling this vector to have a mean of one (Gelman *et al*. 2021, p. 148).

### Diet similarity using model-based ordination

To quantify the main patterns and similarities in the diets of cod and flounder, we used a model-based unconstrained ordination approach (Warton *et al*. 2012, Hui *et al*. 2015). Model-based ordination explicitly models the mean-variance relationship, which is typical in ecological data and if not accounted for, can lead to misleading distance-based ordination plots (Hui *et al*. 2015). We used a generalized linear latent variable model (GLLVM), which fits multivariate data using a factor analytic approach with a set of latent variables that can be interpreted as unconstrained ordination axes. The multivariate data in this case are the relative prey weight (prey-to-predator weight ratio) of the 15 prey groups depicted in Fig. 2a. These groups follow Haase *et al*. (2020), and encompass all true prey items found in the stomachs. We calculated the weighted mean weight of each prey group across space and time (year, quarter) by 2 cm size bins for cod and flounder separately for fish sizes larger or equal to 10 cm and smaller than 50. The averaging by 2 cm size group was because the model could not capture the left skew of the residuals when done on 1 cm groups (see Fig. S3 for the final QQ-plot). After removing size groups with fewer than three predators, the sample sizes across bins ranged between 17–508, with a mean of 275 and a median of 287. This resulted in a 35×15 matrix of stomachs, where the 35 rows correspond to average relative prey weights for a 2 cm class of cod or flounder. We then fit a GLLVM with two latent variables and no covariates, and a Tweedie distribution with a log link function because of the continuous positive response variable and presence of zeroes. The model can be written as:

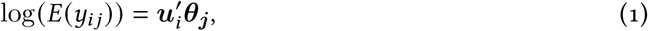

**Figure 2:**
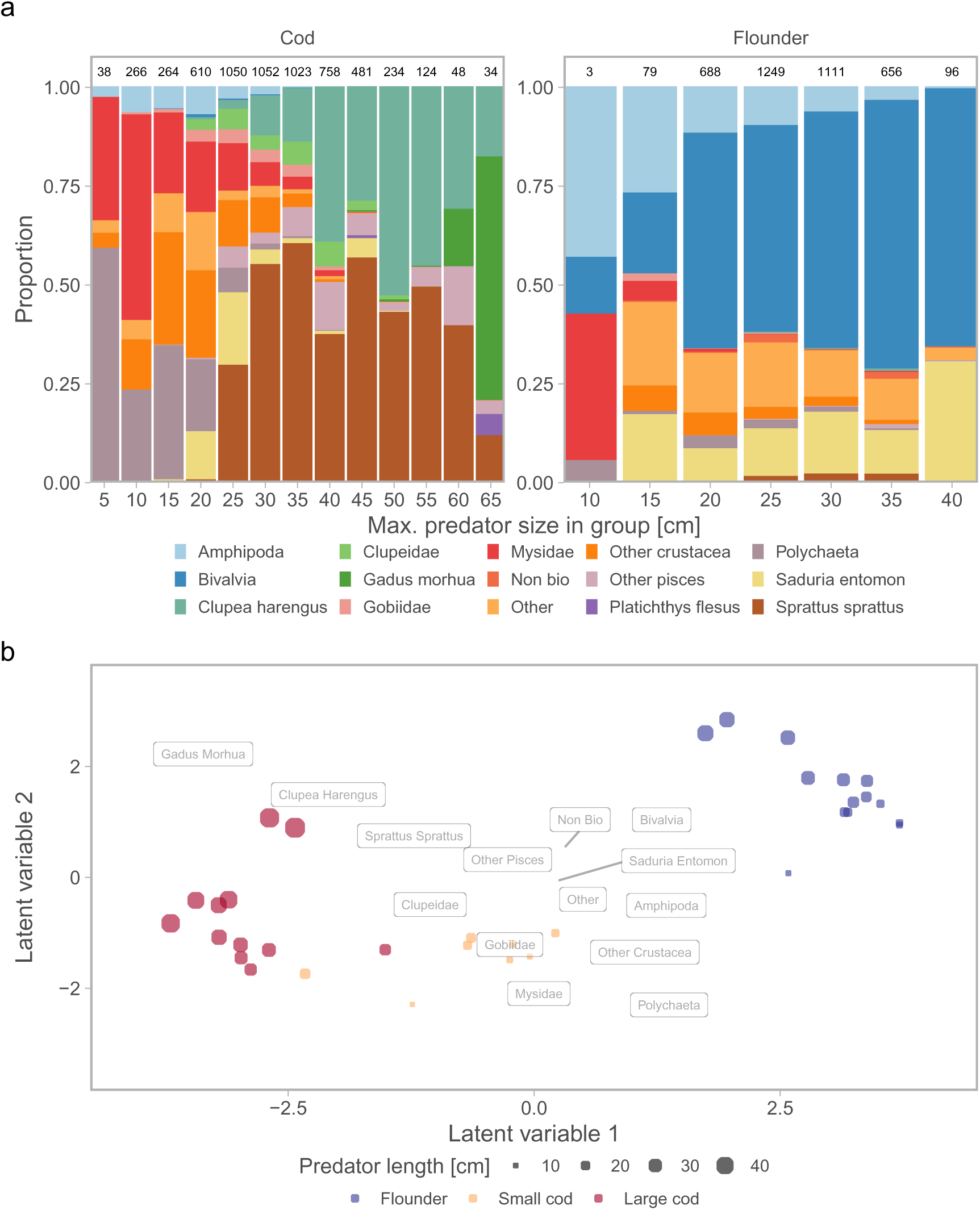
Diet over ontogeny and diet overlap between cod and flounder. Panel a depicts diet as weight proportions by 5 cm size groups of the predator and prey groups (the size is the maximum in that group). Diet proportions are based on weighted means, where the sample weights are the mean-centred ratio of the number of fish caught of the predator group to the number of fish sampled in the associated haul. The number above each bar corresponds to the sample size in each group. Panel b depicts the unconstrained ordination constructed from the two latent variables in the multivariate generalized linear latent variable models (GLLVM), where each point is a 2 cm predator×size group, colors indicate predator groups (flounder in blue, small cod in yellow and large cod in red), and point size (radius) corresponds to the predator length in 2 cm groups. Only predators larger or equal than 10 cm and predators smaller than 50 cm where included in the GLLVM model.

where *E* is the expected value, *y*_*i* *j*_ is the average relative prey weight of prey *j* in predator-at-size *i*, ***θ_j_*** are latent variable loadings (coefficients related to the latent variables ***u***_*i*_) that provide coordinates for average stomach contents in the ordination plot, which allows for a graphical representation of stomachs that are similar in terms of their prey composition. We visualized the ordination by species and size. Specifically, small cod were defined as being ≥ 10 and *<* 25 cm, and large cod were defined as being ≥ 25 and *<* 50 cm. The separation between small and large cod was based on the proportion of pelagic food in the diet, where the diets of cod larger than 25 cm are made up by sprat and herring by more than 50% (Fig. 2a). These groups are also used in subsequent analysis. We fit this model in R version 4.3.2 (R Core Team 2023) with the R package gllvm (Niku *et al*. 2019, 2023) (version 1.4.3), which maximizes the log-likelihood of a TMB (Template Model Builder) (Kristensen *et al*. 2016) model with random effects integrated over using the Laplace approximation. We evaluated the model fit using QQ-plots based on randomized quantile residuals (Fig. S3).

### Schoener’s diet overlap index

Next we tested if diet similarity varies with biomass density of potential competitors, cod, and flounder. The hypothesis here is that if potentially competing species occur in high densities at the same place at the same time, there would be a higher diet separation (resource partitioning) to reduce competition (Macarthur and Levins 1967). To this end, we calculated the Schoener’s overlap index (Schoener 1968, see also e.g. Barnes *et al*. (2018)), and estimated the effects of pooled competitor biomass on the diet overlap.

Schoener’s overlap is a symmetric measure of niche equivalency and varies between 0 (no overlap) and 1 (complete overlap), where values exceeding 0.6 are typically viewed as important (Zaret and Rand 1971, Wallace 1981, Christensen *et al*. 2008). It is calculated as:

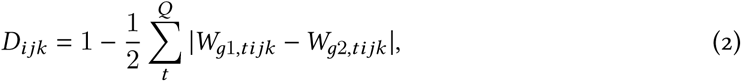

where *D*_*i* *jk*_ is the overlap index by year *i*, quarter *j*, and ICES rectangle *k*. *W*_*g*1,*ti* *j*_ and *W*_*g*2,*ti* *j*_ are the proportions of prey taxa *t* by weight in the stomachs of predator group 1 and 2, where predator group is flounder, small cod, and large cod. The variable *Q* is the total number of prey taxa observed. We calculated weighted mean prey weights by year *i*, quarter *j* and ICES rectangle *k* to balance getting representative diets of the predator pairs and a large enough sample size to ensure a contrast in both diet overlap and competitor biomass density in the subsequent analysis described below. We verified that this resulted in large enough sample sizes within each spatiotemporal unit to acquire representative diets on average (Fig. S4). We used the same prey taxa groups as in the GLLVM model and the same filtering, yielding 259 overlap values. This analysis represents the intermediate spatial scale, i.e., at the ICES rectangle level, which is 1 degree of longitude by 0.5 degrees of latitude (see Fig. 1).

To model the mean overlap, and the effects of biomass on overlap, we fit a beta GLM, because overlap is a proportion. We replaced exact zeroes of overlap with the lowest positive value (2.7% of rows changed from 0 to 0.0003), because the beta distribution does not allow exact zeroes. We opted to do this instead of fitting a zero-inflated beta model because not all predator-pairs had zeroes and we used overlap pair (combination of flounder, small cod and large cod) as a predictor, and because not all bootstrapped data sets (described below) necessarily include the few zero observations. To evaluate if the pooled biomass density (kg/km^2^) of cod and flounder from the trawl surveys affects the diet overlap index, we included the natural logarithm of the mean biomass density across small cod, large cod, and flounder for the same subset of ICES rectangles and years for which we had overlap data. The biomass density variable was scaled by subtracting the mean and dividing by the standard deviation. We included mean predator biomass density as a predictor for the mean of the beta distribution and let it interact with overlap pair to allow biomass density to have different effects on each predator overlap combination. The model can be written as:

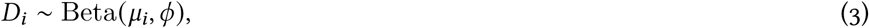

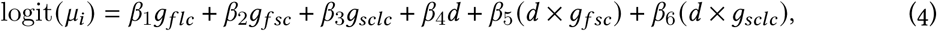

where *D*_*i*_ is the overlap index, *μ*_*i*_ and *ϕ*_*i*_ represent the mean and the precision of the beta distribution, respectively. The symbol *g* is an indicator variable that denotes overlap pair, such that *β*_1−3_ are the mean overlap indices for pairs of flounder and large cod (subscript _*f* *lc*_), flounder and small cod (subscript _*f* *sc*_), and small cod and large cod (subscript _*sclc*_), respectively. The parameter *β*_4_ is the slope of biomass density (*d*) for the overlap group of flounder and large cod, *β*_5_ is the difference in biomass density slope relative to *β*_4_ for the group flounder and small cod, and *β*_6_ is the difference biomass density slope relative to *β*_4_ for the group small cod and large cod. We fit the model using the R package glmmTMB (Brooks *et al*. 2017) (version 1.1.10), and evaluated model fit by visually inspecting scaled, simulation-based residuals calculated using the R package DHARMa (Hartig 2022) (Fig. S6).

We used weighted means to calculate Schoener’s overlap metric, which corrected for diet samples coming from large or small catches, but we also needed to correct for these weighted means being calculated in spatiotemporal units with different numbers of samples. To account for this, we acquired uncertainty estimates from a bootstrap procedure. The procedure was as follows. For each of 10,000 bootstrap samples: (1) sample data with replacement, (2) calculate the diet overlap index, and (3) fit the beta model and extract coefficients. Uncertainty on intercepts (mean overlap) and slopes (effect of biomass on overlap) were characterized by the 5^th^ and 95^th^ percentiles of the distribution of parameter estimates. Bootstrapping was done with the R package boot (Davison and Hinkley 1997, Canty and Ripley 2022).

### Spatiotemporal food competition models

Both the ordination and the Schoener’s overlap index require some spatial aggregation, and hence do not address local-scale variability and effects. However, competitive interactions may also vary on more local scales than the rather arbitrary ICES rectangle (Schoener’s overlap analysis) or the entire domain (GLLVM ordination). However, one can imagine a scenario where the food consumed (total or particular prey) rather than the prey composition varies with predator density, and that biomass of prey (total or particular prey) in predator stomachs is more closely related to competition and fitness than prey composition. Therefore, in our last analysis, we tested if the relative prey weight in the diet of individual predators was affected by the local biomass of potential competitors. We did this by fitting spatiotemporal GLMMs to relative prey weights of individual predators (see Fig. S2 for a plot of the data). As in Grüss *et al*. (2023), we chose to use individual predators as the sampling unit rather than haul-averaged diets even though stomachs within a haul may not be independent. This is because diets vary considerably with predator size and we could include predator size as a covariate with individual fish. We fit models to flounder, small cod, and large cod separately.

### Response variables

We used three different response variables: relative weight of (1) the benthic isopod *Saduria enotomon*, (2) all benthic prey, and (3) total prey. Benthic prey is here defined as the sum of all prey except pelagic fish (sprat, herring and Clupeidae, see Fig. 2a). Total prey weight was only fit to the large cod group since nearly all prey are benthic for the other predator groups. The rationale for this model setup was as follows: we looked specifically at *Saduria* because it is a prey that is less common in cod stomachs in the 2000s compared to the 1960s–70s (Kulatska *et al*. 2019, Haase *et al*. 2020, Neuenfeldt *et al*. 2020), and this reduction has been linked to the decreased growth of cod (Neuenfeldt *et al*. 2020). Moreover, it is one of the few prey cod and flounder have in common in relatively large numbers (*>* 10% weight-proportion for a 20 cm cod and 10–30% of flounder diets depending on its size, see Fig. 2a). Hence, the *Saduria* models were to test if there is competition for a limited *Saduria* resource. With the benthic prey weight models, we aimed to evaluate if there was evidence for competition for benthic prey more broadly. Competition for *Saduria* in particular and benthos in general has been hypothesised to occur due to the combined effect of the range contraction of cod (Orio *et al*. 2019, 2020) and the expansion of hypoxic areas (Casini *et al*. 2016b, Haase *et al*. 2020). We fit the total prey model for large cod to understand if large cod are able to compensate with pelagic food to keep total food contents constant, should there be competition for benthic food.

Covariates

We matched monthly predicted values of dissolved oxygen at the sea floor with individual-level stomach data. These stem from the biogeochemical model ERGOM (Ecological Regional Ocean Model, https://ergom.net/), using the version in Neumann *et al*. (2015), downloaded from the Copernicus Marine Service Information (CMEMS 2023). As a sensitivity analysis, we also fitted the models to oxygen data acquired from CTD samples (1 m from the sea floor) taken at or near the location of the trawl hauls where stomachs were collected. Using measured data as a covariate would have been the first choice. However, since these measured data come with other limitations, such as variability in data quality and quantity, as well as uncertainty with respect to the distance from the sea bottom, we opted for the modelled oxygen data as the main choice, followed by a sensitivity analysis where we compared the estimates of modelled and measured oxygen (Fig. S13). We also used depth as a covariate, since it greatly influences benthic communities in the Baltic Sea (Karlson *et al*. 2002). Depth was matched to the stomach data using raster files from the EMODnet Bathymetry project (https://emodnet.ec.europa.eu/en/bathymetry) (EMODnet Bathymetry Consortium 2022).

We followed the approach in Lindmark *et al*. (2023), and acquired predicted values of *Saduria* biomass density from the habitat distribution model in Gogina *et al*. (2020), applied with modelled hydrographical data from the regional coupled ocean biogeochemical model ERGOM (Neumann *et al*. 2015, 2021). The *Saduria* model was trained to the time period 1981–2019 and predicted for the time period 2015–2020. Therefore, this variable represents spatial but not temporal variation in *Saduria* biomass density (Fig. S11). We acknowledge this is not ideal and it would be favourable to use real observations instead. However, we deemed it was not possible to accurately link *Saduria* samples from e.g., national monitoring programs or trawl surveys to diet data, and therefore chose to use model predicted values. Moreover, a recent study found that this modelled density correlated positively with *Saduria* in cod stomachs over a 30-year time period (Lindmark *et al*. 2025). We used the natural logarithm of haul-level biomass densities (kg/km^2^) of cod and flounder from the same BITS survey that collected stomach data as covariates to evaluate the effects of inter- and intraspecific food competition. We replaced zeroes in the covariate with half the minimum observed value for that predator. We included interactions between biomass density of flounder and *Saduria*, and small cod and *Saduria* for the models fitted to relative weights of *Saduria*. For the benthic and total prey weight models, we included interactions between flounder and oxygen and small cod and oxygen. This served to explore the hypothesis that the effects of competitor biomass on stomach contents are stronger when prey are scarce. Oxygen is known to affect benthic prey biomass and in particular their depth distribution (Karlson *et al*. 2002, Ojaveer *et al*. 2010, Gogina *et al*. 2016). Hence, the oxygen variable relates both to the direct effect of dissolved oxygen on food consumption (via metabolic rate), and as a proxy related to benthic prey abundance. Lastly, we included individual predator length as a covariate, because even within predator-size groups, size affects the relative importance of benthic and pelagic diets in cod mainly. All variables were scaled by subtracting the mean and dividing by the standard deviation. None of the covariates included in a single model had absolute correlation coefficients *>* 0.5 with each other.

### Model description

The spatiotemporal GLMMs were assumed to follow a Tweedie distribution because of the presence of zeroes and the positive continuous values of relative prey weights. Despite the response always being *<* 1 (prey weigh less than predators), we chose a Tweedie observation model over a beta hurdle model (a Bernoulli distribution for zero vs. non-zero and a beta distribution for the positive ratio) after inspecting residuals. The spatiotemporal models can be written as:

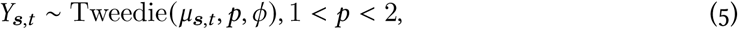

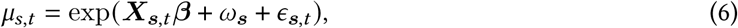

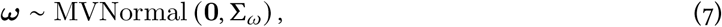

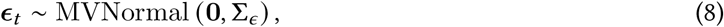

where *Y*_*s*,*t*_ represents the relative prey weight (*Saduria*, benthic or all prey) of an individual predator in space ***s*** (a vector of two UTM zone 33 coordinates) and time *t* (year), *μ* represents the mean relative prey weight, *p* represents the Tweedie power parameter, and *ϕ* represents the Tweedie dispersion parameter. The symbol ***X*** represents the design matrix and ***β*** represents a vector of fixed effect coefficients. The covariates include year and quarter (as categorical variables), individual predator length, depth, oxygen, and biomass density of small cod, large cod, flounder and *Saduria*, including some interactions (see section *Covariates* for reasoning on which interactions to consider). Table S1 shows the pseudo R code for these models.

The spatial and spatiotemporal random effects *ω****_s_*** and *∈****_s_***_,*t*_ capture spatially correlated latent effects that are constant through time and that vary through time, respectively. For example, here spatially constant latent effects might include sediment type, which is largely constant through time, and spatiotemporal latent effects might include plankton productivity, which changes through time. These random effects are assumed drawn from Gaussian Markov random fields (GMRFs) with covariance matrices **Σ**_*ω*_ and **Σ**_*∈*_ constrained by Matérn covariance functions (Rue *et al*. 2009, Lindgren *et al*. 2011). The Stochastic Partial Differential Equation (SPDE) approach (Lindgren *et al*. 2011), which links Gaussian random fields with GMRFs, requires piece-wise linear basis functions defined by a triangulation, commonly referred to as a “mesh”. We formed this mesh using triangles with a minimum distance between vertices of 5 km — i.e., a “cutof” of 5 km and all other arguments in the function fm_rcdt_2d_inla() from the fmesher R package (Lindgren 2023) at their defaults (Figs. S7–S9). We ensured the estimated range at which spatial correlation is present was, as a minimum, approximately twice the minimum distance between vertices (Fig. S12). For each model, we fit four different random effect structures: no random effects, spatial random effects, spatiotemporal random effects, and spatial and spatiotemporal random effects. We compared these structures using the marginal Akaike Information Criterion (AIC) (Akaike 1974), and selected the random effect structure that had the lowest AIC (Table 1).

**Table 1:** ΔAIC (AIC for the model relative to the model with the lowest AIC) for all spatiotemporal GLLMs fitted to stomach content data.

| Response variable | Group | $\Delta$ AIC | | | |
| --- | --- | --- | --- | --- | --- |
|  |  | Non-spatial | Spatial | Spatiotemporal | Spatial & Spatiotemporal |
| Benthic prey | Large cod | 463 | 240 | 0 | 0.353 |
| Benthic prey | Small cod | 941 | 249 | 0 | 2 |
| Benthic prey | Flounder | 312 | 78.8 | 12.5 | 0 |
| *Saduria* | Large cod | 335 | 27.6 | 14.4 | 0 |
| *Saduria* | Small cod | 615 | 58.4 | 18.9 | 0 |
| *Saduria* | Flounder | 567 | 88.6 | 35.7 | 0 |
| Total prey | Large cod | 373 | 156 | 0.35 | 0 |

Our sample weights, which correct for diet samples coming from hauls with different total catches, where multiplied with the corresponding log likelihood for each observation. A comparison of parameter estimates from weighted and unweighted models can be found in Fig. S14.

### Model fitting

We fit the spatiotemporal models with the R package sdmTMB (Anderson *et al*. 2024), version 0.6.0.9034. sdmTMB uses automatic differentiation and the Laplace approximation from the R package TMB (Kristensen *et al*. 2016) and sparse matrix structures to set up the SPDE-based GMRFs from the R package fmesher (Lindgren 2023). Estimation is done via maximum marginal likelihood. We confirmed that the non-linear optimization was consistent with convergence by checking that the Hessian matrix was positive definite and the maximum absolute log-likelihood gradient with respect to fixed effects was *<* 0.001. We evaluated consistency of the model with the data by calculating randomized quantile residuals (Dunn and Smyth 1996) with fixed effects held at their maximum likelihood estimates and random effects sampled via a single draw from Markov chain Monte Carlo (MCMC) (Thygesen *et al*. 2017) using Stan (Carpenter *et al*. 2017, Stan Development Team 2024) via tmbstan (Monnahan and Kristensen 2018) (Fig. S10).

## Results

### Diet similarity

The diet of cod changes with size (Fig. 2a). In terms of prey weight, the smallest cod (5 cm) feed predominantly on polychaetes. As cod further grow in size, crustaceans, of which Mysidae is the largest group, start to dominate their diet. *Saduria* and sprat become a part of the cod diet at around 15–20 cm, and above 30 cm, sprat and herring together constitute more than half of the prey biomass. The proportion of fish in the diet of cod continues to increase with size. Cod and flounder emerge in the cod diet at around 55–60 cm.

The ontogenetic diet shift in flounder is more subtle. Flounder diets do not change dramatically over ontogeny (note the dominance of mysids in the smallest flounder group is based on a small sample size). Overall, their diet is dominated by bivalves, *Saduria*, and other crustaceans (Fig. 2a).

The ordination plots constructed from the first two latent variables revealed a clear clustering by species and for cod also by size (Fig. 2b). Flounder occupied a distinct position on the horizontal axis, associated with benthic prey such as amphipods, bivalvia, and polychaetes. Larger cod were more strongly associated with fish prey including clupeids (*Clupea harengus* and *Sprattus sprattus*).

### Effects of competitor biomass on diet overlap

Schoener’s dietary overlap metrics varied across units (ICES rectangle, year, and quarter combinations between 0.0003 and 0.95 (Fig. 3a). The average overlap was low; 0.1 [95% confidence interval based on percentiles of the bootstrapped estimates: 0.09 0.12] for flounder and large cod, 0.19 [0.15, 0.19] for flounder and small cod, and 0.24 [0.18, 0.24] for small cod and large cod (Fig. 3a). This can be compared to 0.6, which is often considered to be ecologically significant (Zaret and Rand 1971, Wallace 1981, Christensen *et al*. 2008). We find varying effects of biomass density of competitors on the mean of the beta distribution (Fig. 3b). For the overlap between small and large cod, the estimate is -0.2 [-0.5, -0.15], for the overlap between flounder and small cod the estimate is 0.05 [-0.14, 0.13], and for flounder and large cod it is 0.14 [0.08, 0.20] (Fig. 3b). To exemplify these effects, for flounder and large cod, the overlap increases from 0.08 to 0.14, and the overlap between small and large cod declines from 0.32 to 0.15 when the biomass density increases from the lowest to the highest value.

**Figure 3:**
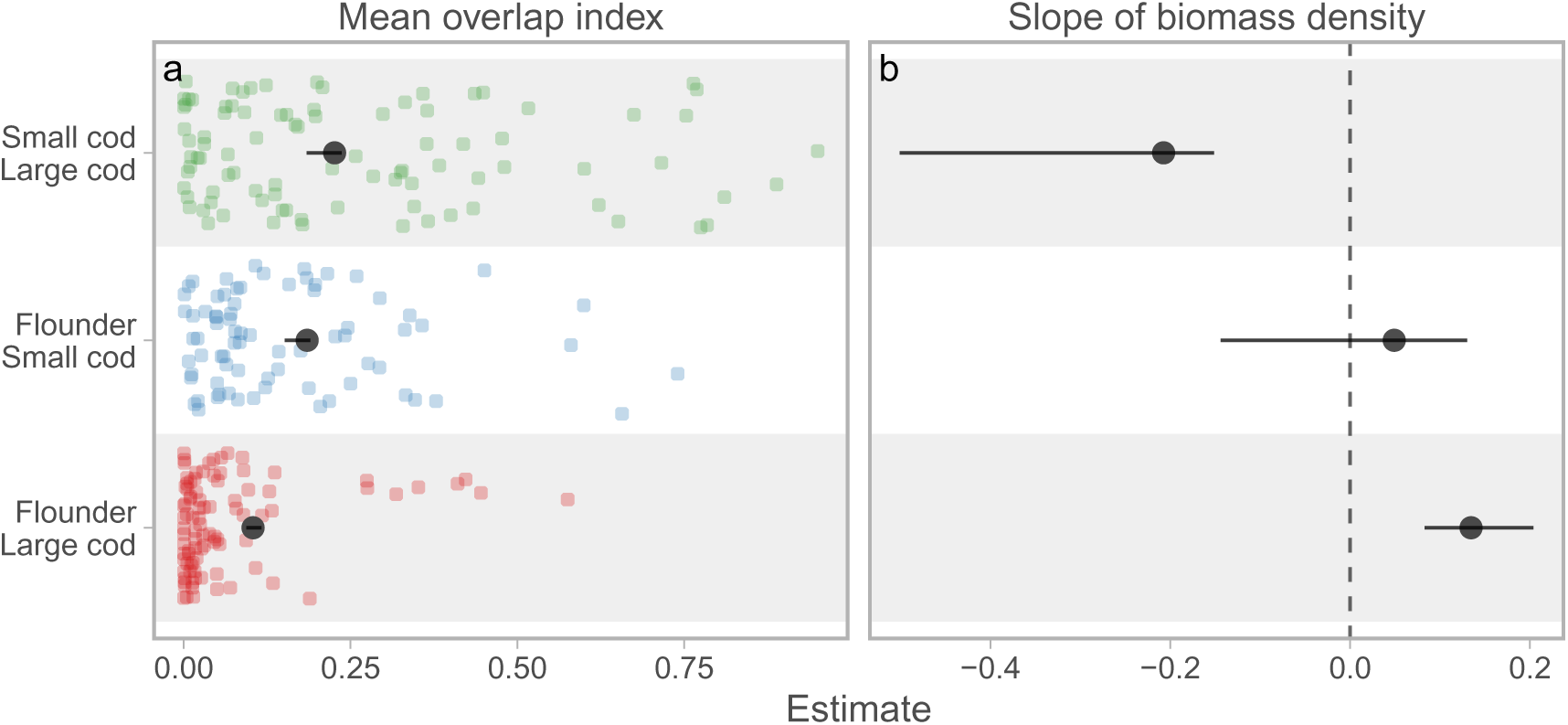
Schoener’s diet overlap index and the effects of biomass density. Panel a depicts the overlap values (x-axis) across all combinations of predator pairs (y-axis, indicated by color), where each point represents an overlap value in a given ICES rectangle, year and quarter. The black points correspond to the average predicted overlap index (intercepts) from the beta GLM. Panel b depicts the slope of average predator biomass density in the same spatiotemporal unit. Horizontal error bars correspond to 95% confidence intervals based on percentiles of the bootstrap distribution.

### Spatiotemporal diet GLLMs

Across all predator and prey groups, the AIC model comparison either favoured models with (1) both spatial and spatiotemporal random fields (all *Saduria* models, the benthic prey model for flounder, and the total prey weight model for large cod) or (2) spatiotemporal random fields (benthic prey model for small and large cod) (Table 1). However, some differences are small (ΔAIC≤2 for spatiotemporal vs. spatial and spatiotemporal models for benthic and total prey models for cod). The spatial and spatiotemporal random effects were larger in magnitude and the spatial clustering was more pronounced due to the high degree of latent variable spatial correlation within the *Saduria* models compared to the benthic and total prey models (Fig. S17–S19).

The relative weight of *Saduria* in stomachs was negatively associated with flounder biomass density in all predator groups (Fig. 4a). The predicted availability of *Saduria* was associated with higher relative weights of *Saduria* in small cod and large cod mainly, but for the latter the 95% confidence interval overlapped 0 (Fig. 4a). The interactions between flounder and predicted availability of *Saduria* overlapped 0 for all predators (Fig. 4a), suggesting the density of *Saduria* in the environment does not matter for the effect of flounder on the relative weights of *Saduria* in stomachs. This can also be seen in the conditional predictions of flounder density for two densities of predicted *Saduria* availability (Fig. 5). However, a limitation here is that the *Saduria* densities are predicted values and have no temporal variation. Relative *Saduria* weights in stomachs were positively associated with the density of small cod in flounder predators. Relative weights of benthic prey were negatively associated with flounder density in flounder predators, but not associated with flounder density in neither small nor large cod (Fig. 4b). The effect of large cod biomass density on the total relative prey weight in large cod was negative.

**Figure 4:**
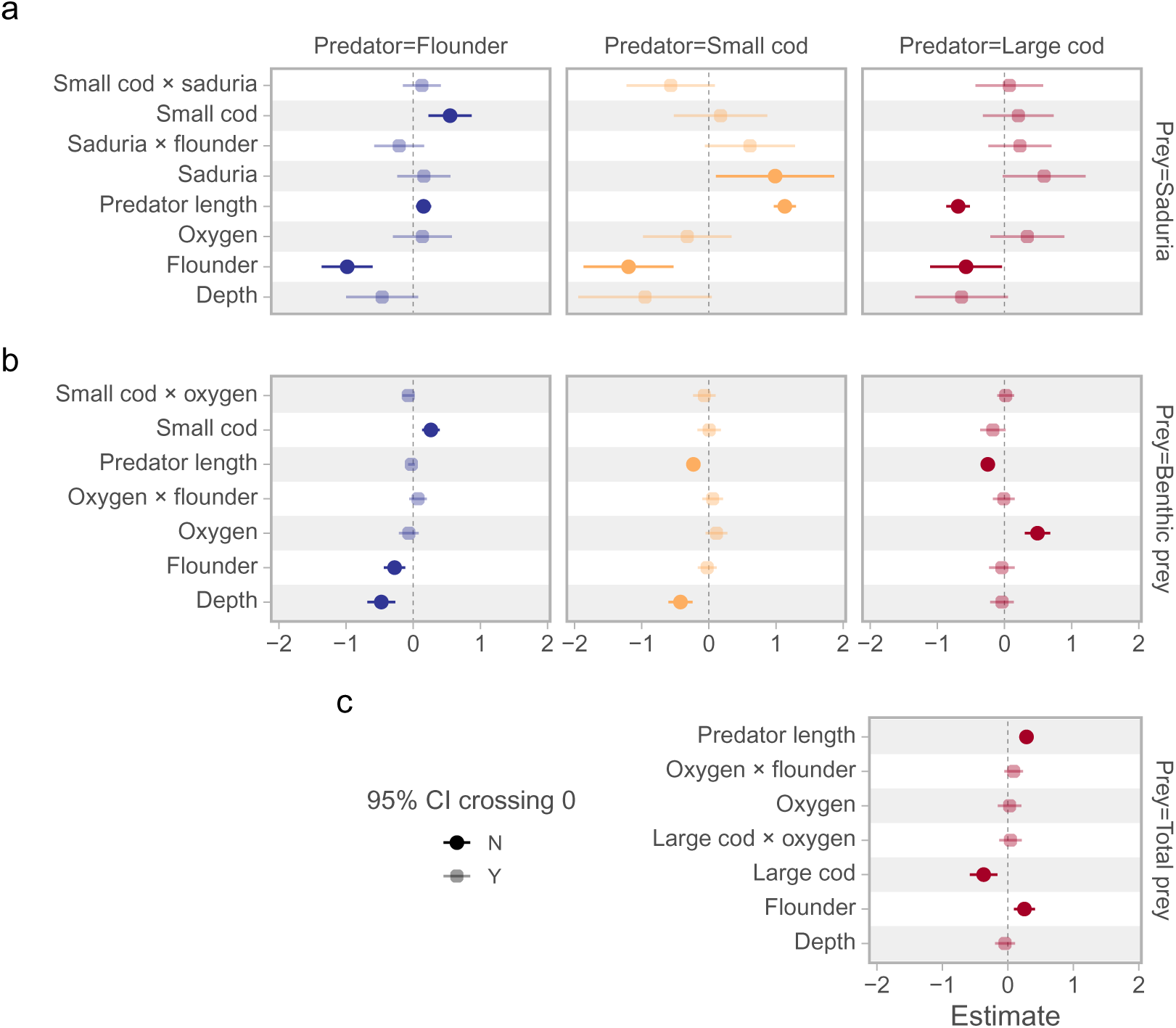
Standardized coefficients from the spatiotemporal generalized linear mixed model fitted to relative prey weights. Predator groups are in columns (also indicated with color), and each row represents a prey group (response variable, a=*Saduria*, b=Benthic prey, and c=Total prey weight). The y-axis shows the predictor variable. The transparency corresponds to the support (yes/no = Y/N) for the effect, such that semi-transparent points show estimates with 95% confidence intervals overlapping 0.

**Figure 5:**
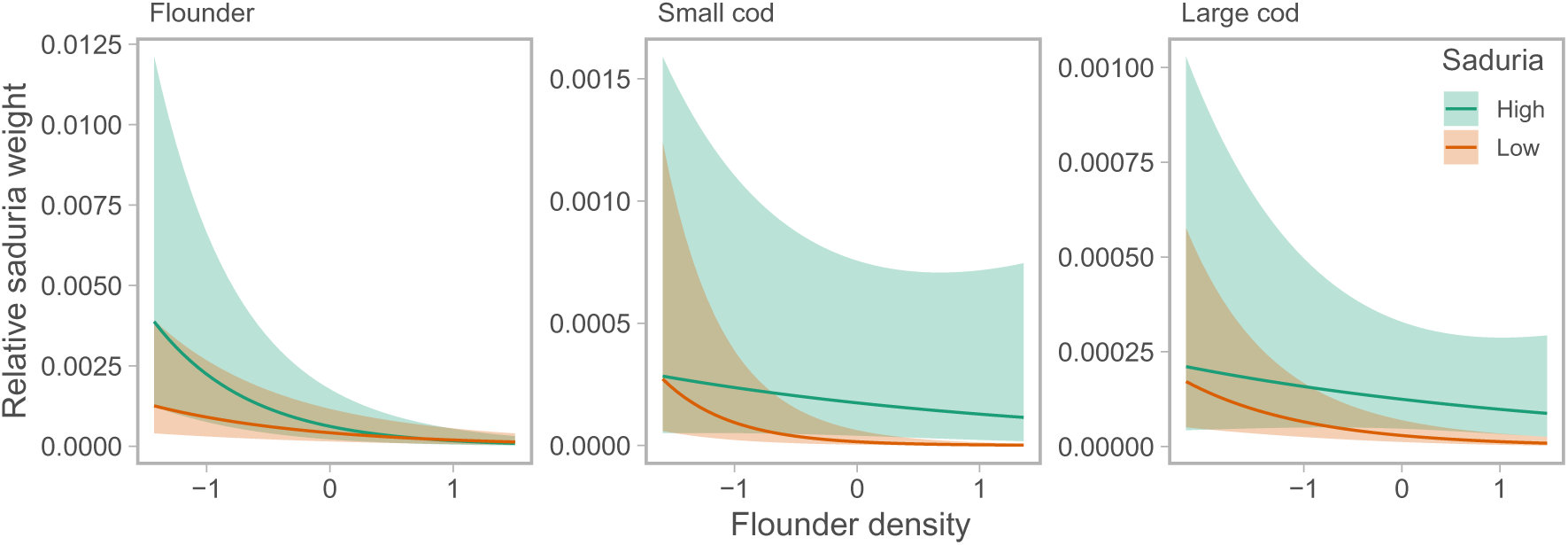
Conditional effect on the response scale of flounder density on the relative weight of *Saduria* in stomachs of flounder (left), large cod (middle) and small cod (right), for two levels of *Saduria*: high in green (95^th^ percentile) and low in orange (5^th^ percentile). All other covariates are held at their mean, quarter is set 1 and year is 2020, and random fields are set to 0. The ribbon corresponds to the 75% confidence interval, to illustrate differences between high and low *Saduria* availability while still illustrating the uncertainty of the prediction. Note the different y-axes.

Depth was negatively related to relative prey weights of *Saduria* and benthic prey in flounder predators, while depth did not correlate with relative benthic or total prey weights in large cod predators. *Saduria* was negatively related to depth in small and large cod, but the 95% confidence interval overlapped 0. The effect of oxygen on relative *Saduria* weights was nearly 0 for flounder, positive for large cod, and negative for small cod predators, but in all cases the 95% confidence intervals overlapped 0 (Fig. 4a). With benthic prey as the response variable, oxygen had a clear positive effect for large cod, but not for flounder or small cod (Fig. 4b). While the absolute oxygen estimates varied depending on whether oxygen measurements came from CTD or modelled oxygen, the sensitivity analysis resulted in sign-agreement in the oxygen estimates for all models except for benthic prey in flounder predators (but both oxygen sources result in oxygen coefficients with confidence intervals overlapping 0); see the oxygen estimates in Fig. S13a. The positive effect of oxygen on benthic prey changed from clearly positive to overlapping with 0 when using observed rather than modelled oxygen in large cod (Fig. S13b).

There was interannual variation in the relative weights of benthic prey and total prey for all predators, but the uncertainty of the prediction made it difficult to detect any trends in the short time series. The effect of quarter was more similar within prey groups than across predators (Fig. S16). For benthic prey and cod, the relative weights tended to be higher in the fourth than the first quarter, and for *Saduria* relative weights were larger in the fourth quarter for small cod and flounder (Fig. S16).

## Discussion

In this study we evaluated if spatiotemporal dietary patterns of cod and flounder in the Baltic Sea are consistent with food competition using statistical modelling of stomach content data at different spatial scales. Overall, we found low dietary overlap between these two predators. Moreover, on a intermediate spatial scale (ICES rectangle, 1 × 0.5 degrees longitude and latitude grid cells, Fig. 1), we did not find support for increased diet differentiation as predator densities increased, as is often expected under resource partitioning to reduce competition and facilitate coexistence (Macarthur and Levins 1967). However, at the local scale (individual stomachs at the trawl haul level), we find that the relative weights of the isopod prey *Saduria entomon* in the diets of flounder, small and large cod decline with increasing flounder densities. The total prey weight in the cod stomachs on the other hand is not affected by flounder density — large cod do not have less benthic prey or total prey, and small cod do not have less benthic prey, when flounder are abundant. Hence it seems the food competition from flounder does not mean less total food in cod stomachs. Rather, our results could mean that cod feed less on *Saduria*, but compensate for this loss with other prey. We also find results consistent with intraspecific food competition; large cod tend to have less prey in stomachs when the density of large cod increases, and the same was found for flounder predators.

We find low diet overlap between cod and flounder, but in itself this does not inform our understanding of competition (Pianka 1976, Schoener 1983). If the resources are limited, a high overlap could indicate a tendency to compete for food, and the intake of the shared resource could decline as the biomass of the potential competitor increases. A low resource overlap could be a result of dietary resource partitioning shaped over evolutionary time scales to avoid competition, but it could also arise as a behavioural response over short time scales as a form of prey switching by a generalist predator. We estimate relatively small negative and positive effects of total predator biomass density on the diet overlap for our three predator groups. This lack of clear negative effect speaks against the hypothesis that high competitor density drives behavioural diet separation.

However, on a local scale (haul-level), we do find signs of what could be resource partitioning and interspecific food competition for *Saduria*, but not declines in the total weight of prey. The findings that cod feed less on *Saduria* when flounder are more abundant, are in line with previous studies on fish documenting behavioural resource partitioning in that it is the more specialist species (in our case, flounder) that successfully interferes with the generalist (cod) (Colwell and Fuentes 1975, Larson 1980). This could indicate competition if the predator replaces *Saduria* with a worse prey in terms of nutrients or energy and fitness is negatively affected by it. It could be that *Saduria* is an especially important prey for cod relative to its weight contribution in the diet. Lindmark *et al*. (2023) found that predicted *Saduria* availability was positively related to body condition (albeit weakly), and while the energy density is lower compared to pelagic prey (Pedersen and Hislop 2001, Neuenfeldt *et al*. 2020), crustaceans, such as *Saduria*, have been considered important dietary components due to their high levels of the carotenoid astaxanthin (Røjbek *et al*. 2012). Speaking against this hypothesis is the observation that between 1963–1974, when the the relative weight proportions of *Saduria* were highest and ranged between 15–25% for cod above 21 cm (Haase *et al*. 2020), somatic growth of cod was the lowest recorded for cod of 25 cm (Mion *et al*. 2021).

While we find negative associations between flounder density and cod feeding on *Saduria* in recent years, this does not necessarily mean that competition is responsible for the long-term decline of *Saduria* in cod stomachs. *Saduria* was more common in the diet of cod prior to the early 1990s (Neuenfeldt *et al*. 2020, Haase *et al*. 2020), and spatial overlap between cod and flounder may have increased over time (Orio *et al*. 2020). However, the timing of these trends is inconsistent; flounder abundance was high in the southern Baltic Sea in the late 1980s (Orio *et al*. 2017), while cod feeding on *Saduria* started to declined primarily after the late 2000s (Lindmark *et al*. 2025). An alternative explanation for why cod feed less on *Saduria* now is that the spatial overlap between cod and *Saduria* has declined (Lindmark *et al*. 2025). It could also be that the *Saduria* population overall has declined, although Svedäng *et al*. (2022) showed that *Saduria* populations in shallow areas have remained relatively stable over the past 30 years. Continued expansion of hypoxic areas may, however, compress *Saduria* habitats (Karlson *et al*. 2002). Unfortunately, there are no data on flounder diets back in time, which otherwise could help disentangle the mechanisms for the decline in *Saduria* in cod diets. Overall, these results highlight the complexity of ecological interactions in ecosystems experiencing rapid environmental and community change, making it difficult to attribute long-term dietary shifts to any single cause.

Our results on interspecific competition for the isopod *Saduria* are not conclusive. In line with competition, the relative prey weight of *Saduria* in both small and large cod are negatively associated with flounder density (although this could also be due to indirect effects, e.g. less *Saduria* in high quality flounder habitats). We find positive effects of predicted *Saduria* availability in the environment and the weight of *Saduria* in predator stomachs (at least for small cod), which suggests they predate on *Saduria* in proportion to its availability, as also found in (Lindmark *et al*. 2025) over a longer time period. However, as *Saduria* availability declines, the effect of flounder ought to be more negative since this implies *Saduria* are less likely to be available in excess. We do not find such interactive effect between *Saduria* and flounder for flounder predators, but we do find it for small and large cod. For cod, the relative weights of *Saduria* in stomachs decline more strongly with increasing flounder densities when *Saduria* availability are lower, although there is considerable uncertainty associated with this effect (Fig. 5).

From a metabolic perspective, feeding activity and food assimilation require oxygen, and the latter becomes more costly as food rations increase. Therefore, a behavioral response to lower oxygen availability is a reduced appetite (Jobling 1997). In line with this theoretical prediction and experimental and in-situ studies supporting it (Chabot and Dutil 1999, Moriarty *et al*. 2020), Brander (2020) suggested that the decline in feeding levels of cod over time could be linked to the negative trends in dissolved oxygen, causing a state of mild hypoxia (Brander 2020). In this study, we for the first time explicitly test if feeding is related to oxygen in Baltic cod by including local dissolved oxygen as a covariate for prey weights in individual-level cod stomachs. We only find statistically clear and positive effects of oxygen in large cod feeding on benthic prey, and no significant effects for any other combination of predator and prey. There is also some uncertainty as to the positive effect of oxygen for benthic prey in large cod because it disappears when using observed rather than modelled oxygen. For modelled oxygen data the positive effect seems fairly large in large cod. The relative weight of benthic prey in the diet increases from 0.0011 to 0.0016 when oxygen increases from 4.8 to 7.6 ml/L — a 45% increase. This range of oxygen values is similar to the lower and upper end of the interquartile range of oxygen conditions cod experienced between 1993–2019 (Lindmark *et al*. 2023). We cannot completely disentangle the direct effects of oxygen affecting fish appetite and the effect of oxygen on prey availability. However, given that oxygen is only statistically related to benthic prey in large cod, and not total prey nor any other predator-prey group combination, and that oxygen strongly affects benthic communities (Karlson *et al*. 2002), it seems likely that the main effect of oxygen is via food availability rather than a direct effect on the appetite.

One reason resource partitioning is not detected in the dietary overlap analysis could be that the spatiotemporal scale used to aggregate the individual diets was too large. When calculating overlap indices, it is important to ensure a large enough sample size per spatial and temporal aggregation unit to characterize a representative diet, given the large individual variation in diets (Barnes *et al*. 2018). However, increasing the scale over which to aggregate individual stomachs comes at the cost of accurately representing the local biotic and abiotic conditions at the time and location of the stomach sample. While our spatiotemporal GLMMs rely on univariate analysis of specific prey or prey groups rather than the whole prey community, a strength is that it avoids this aggregation over arbitrary spatial units. This is a potential reason why we can detect signs of resource partitioning, if part of the variation within aggregation units can be attributed to variation in environmental conditions within units that drive changes in stomach contents.

Are the inconsistent signs of food competition among predator groups, and the ability to switch prey, a general pattern in marine fish communities in coastal and continental shelf ecosystems? It has been hypothesised that the high productivity and often generalist behaviour of many fish species result in low food competition (Sherman 1991, Link 2002). There are numerous examples in the literature supporting and rejecting this, but three well-studied cases illustrate that there might be signals of food competition in large aquatic systems. First, the crash of the Peruvian anchovy, one of the most heavily exploited fish stocks in the world, led to an increase in its competitor Peruvian Pacific sardine, but only partially, as the upwelling system is largely driven by environmental fluctuations (De Vries and Pearcy 1982). Second, the collapse of the Californian sardine in the late 1940s was followed by an increase in Northern anchovy, but not until a decade later. There were no signs of density-dependent body growth, and analysis of sediment cores suggests there have been periods with high abundance of both species (MacCall 1986, Baumgartner *et al*. 1992, Jennings and Kaiser 1998). Last, the exceptionally strong year classes of gadoid species (codfishes) in the North Sea in the late 1960s — referred to as the “gadoid outburst” — could not convincingly be linked to the decline in Atlantic mackerel and North Sea herring stocks (Cushing 1984, Hislop 1996), as would otherwise be natural to assume given that they compete with juvenile gadoids for zooplankton. Furthermore, Ursin (1982) noted that despite the 5-fold increase in biomass of haddock, whiting, and Norway pout in 1968–69 relative to previous years, which is likely to have doubled the stock of demersal fish for years, no negative effect on growth in demersal life stages of gadoid fish was found (Ursin 1982, Jones 1983). Despite the extreme contrasts in biomass in these examples, the inconclusive results seem emblematic in temperate marine fish communities, possibly due to large influences of environmental drivers and an overall ability to find prey (Ursin 1982).

In conclusion, our approach to quantifying competition for food leveraging local-scale diet and biomass data illustrates the importance of scale and type of indicators when assessing competition. For instance, food competition is expected to occur if predators with similar diets co-occur in space, and the diet similarity is expected to decline as predator densities increase (resource partitioning). In our system, we find low dietary overlap, and no overall negative effect of predator density on the overlap at an aggregated scale. However, when modelling local-scale stomach contents of specific shared prey, rather than spatially aggregated diet similarities, we find negative effects of flounder densities on the weights of *Saduria* in stomachs of both small and large cod, but the total prey weights in cod stomachs are not affected by flounder. This could mean that cod, a generalist predator, is able to adapt its foraging in response to prey and predators’ densities, although the energetic effects of this switch are still unknown. We also find results consistent with intraspecific competition, because the relative weights of benthic prey in flounder, and total prey weights in large cod, tend to be lower when the biomass density is higher. We believe that using individual-level data with local-scale covariates in spatiotemporal models, as done here, can improve our understanding of competitive interactions across scales, which is important for developing ecosystem-based management of wild populations.

## Acknowledgements

We are grateful to everyone involved in data collection (at the Swedish University of Agricultural Sciences, Department of Aquatic Resources, Sweden) and stomach analysis (at the National Marine Fisheries Research Institute in Gdynia, Poland); Barbara Bland, Christina Petterson, and Olov Lövgren for help with acquiring CTD data from the Baltic International Trawl Survey; and Hagen Radtke and Ivan Kuznetsov for assistance in acquiring predictions of *S. entomon* densities. We thank P.A. English and two anonymous reviewers for comments that improved this paper.

## Author contributions

Conceptualization (ML, VB, AE, MS, MC), Data curation (ML, MG, MC), Methodology (ML, FM, SA, MG, VB, MS, MO, AE, MC), Formal analysis (ML, FM, SA), Investigation (ML, FM, SA), Visualization (ML, SA), Software (SA), Funding acquisition (MC, AE, MS, VB), Writing – original draft (ML), Writing – review & editing (ML, FM, SA, MG, VB, MS, MO, AE, MC).

## Data Availability Statement

Code and data to reproduce the results are available on GitHub (https://github.com/maxlindmark/cod-interactions) and will be deposited on Zenodo.

