## Supporting Information for "Quantifying competition between two demersal fish species from spatiotemporal stomach content data"

### S1. Diet data

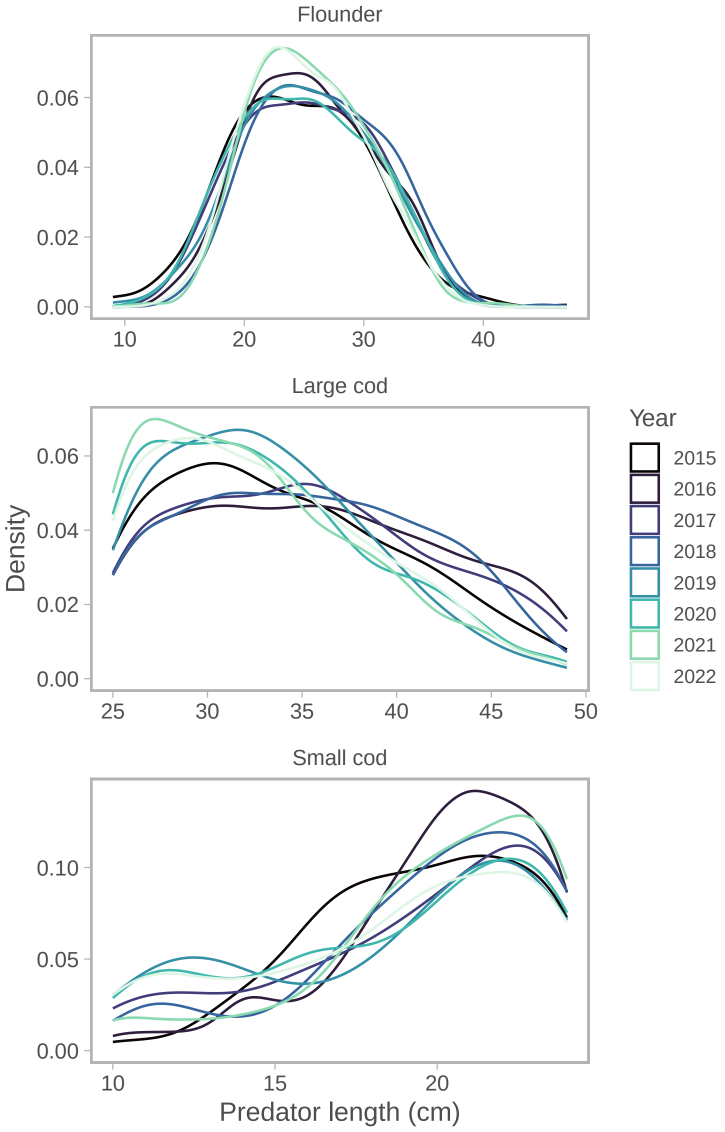


Figure S1: Predator length distributions, by size group and year (color).

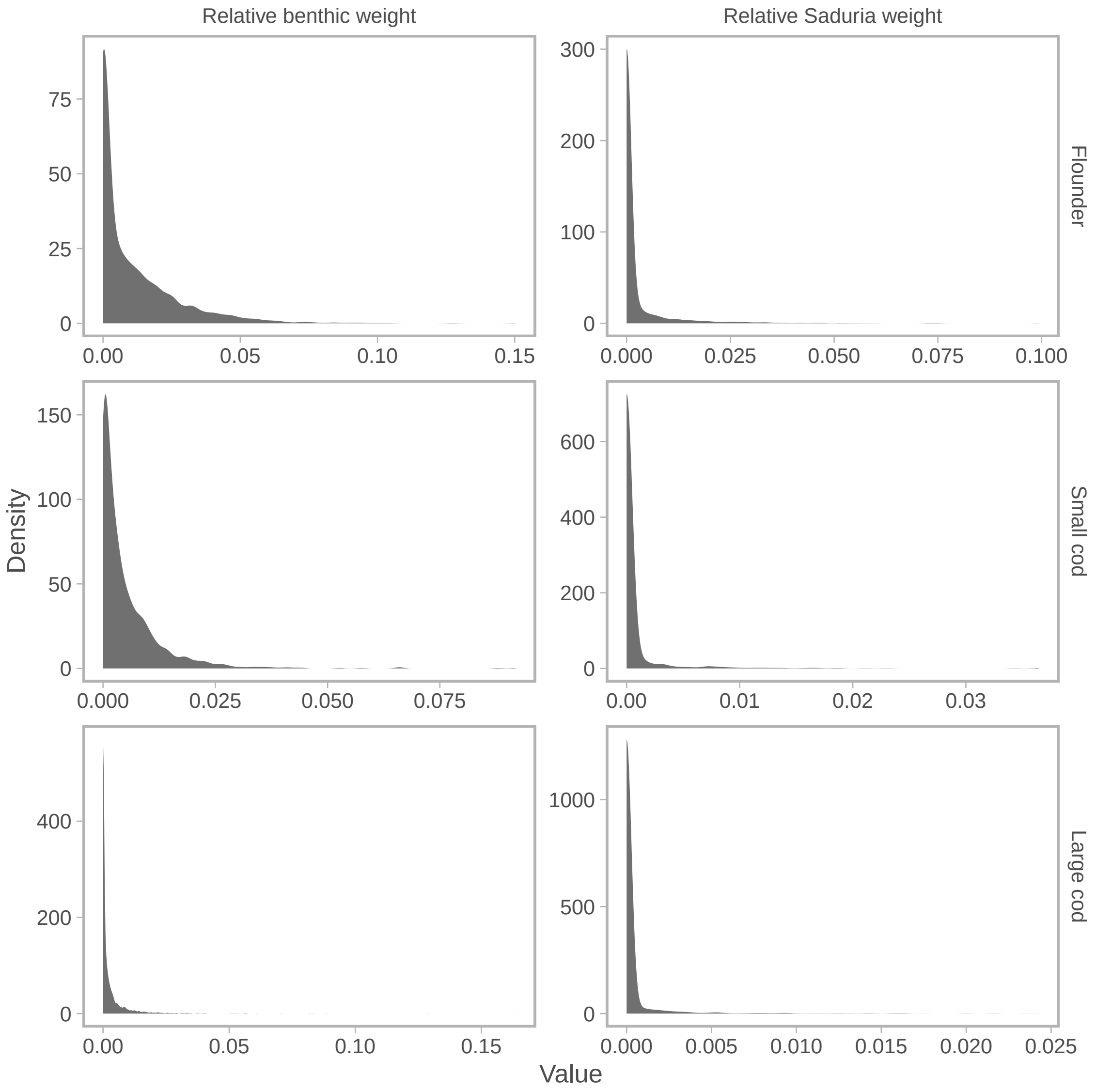


Figure S2: Distribution of response variables (relative prey weights) of benthic prey (left) and *Saduria* (right), for flounder (top), small cod (middle) and large cod (bottom).

### S2. Fit of multivariate diet models for ordination (GLLVM)

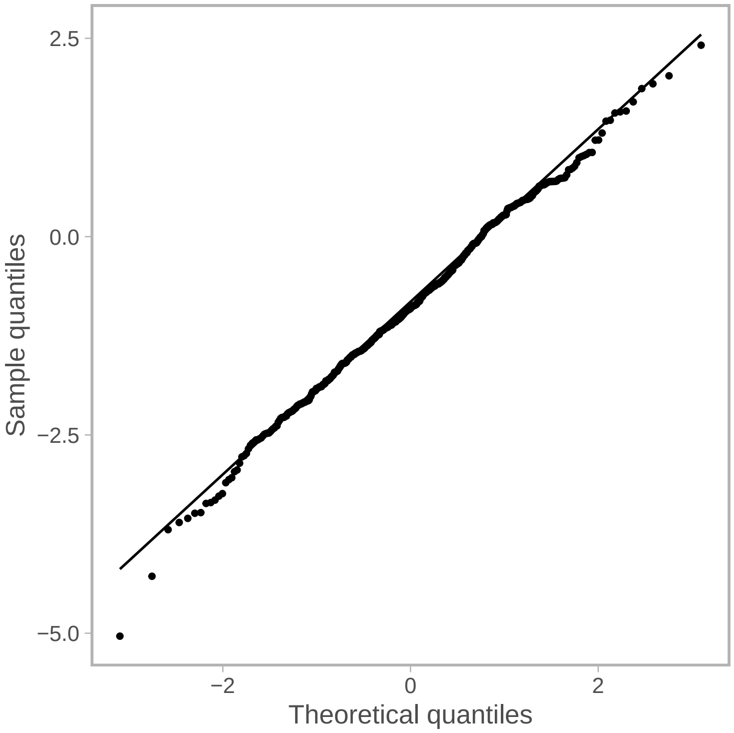


Figure S3: Randomized quantile residuals from the GLLVM model.

### S3. Beta model fitted to diet overlap

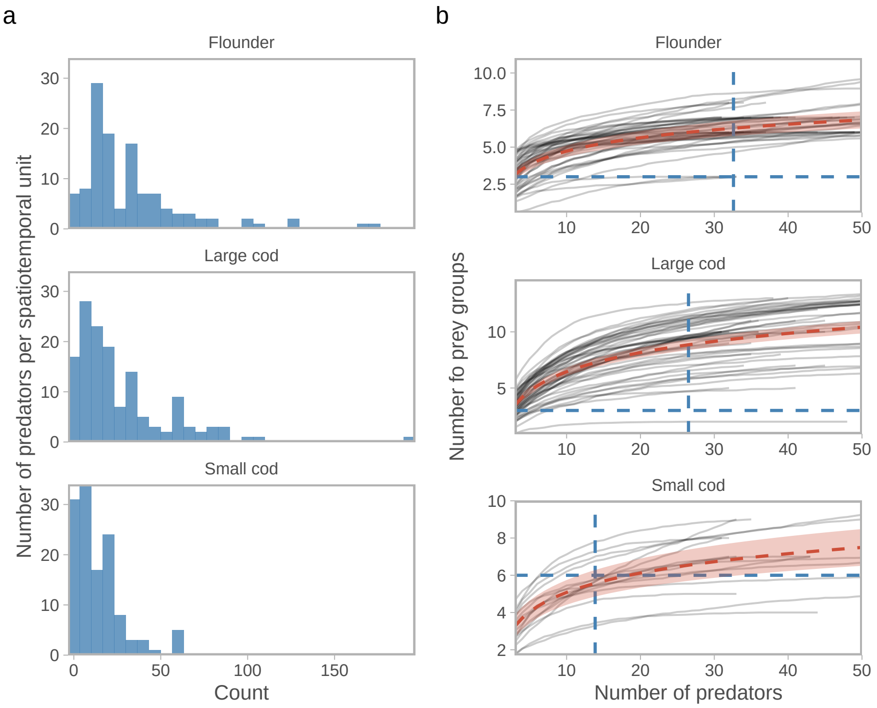


Figure S4: Distribution of sample sizes in spatiotemporal units (combination of year, quarter and ICES rectangle) within which stomachs are aggregated for diet overlap calculations (panel a), and accumulation curves of prey groups in Fig. 2 (panel b), calculated using the function poolaccum() in the vegan package (Oksanen *et al.*, 2022). In panel b, each grey line corresponds to an accumulation curves of prey groups in a single unit. Only groups with more than 30 unique predators are included. The red line corresponds to the global prediction from a linear mixed model fitted to accumulation data points (y ~log(x) * predator_group), with intercepts and slopes varying by unit, fitted with glmmTMB (Brooks *et al.*, 2017). The red area corresponds to the 95% confidence interval. Blue vertical lines correspond to the average number of unique predators across all units, and blue horizontal lines correspond to the number of prey groups each predator group has that contributes to more than 5% of the diet in weight (all stomachs pooled). If blue lines intersect at or below the red line, it indicates that the average number of predators in each unit on average captures equal or more than than the number of unique prey groups that make up more than 5% by weight.

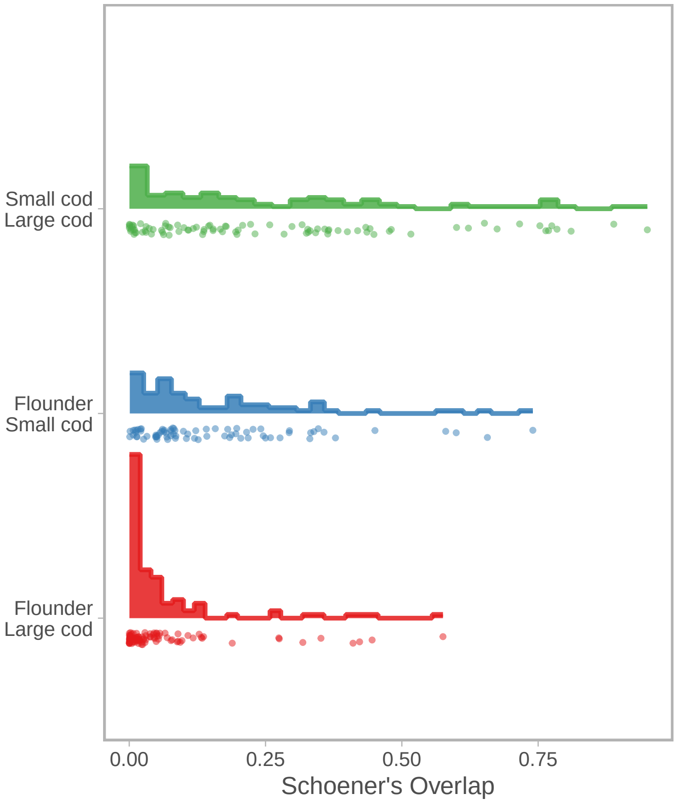


Figure S5: Histograms of overlap values used in beta regression relating overlap to pooled biomass density of competitors. Dots represent the jittered values underlying the histograms.

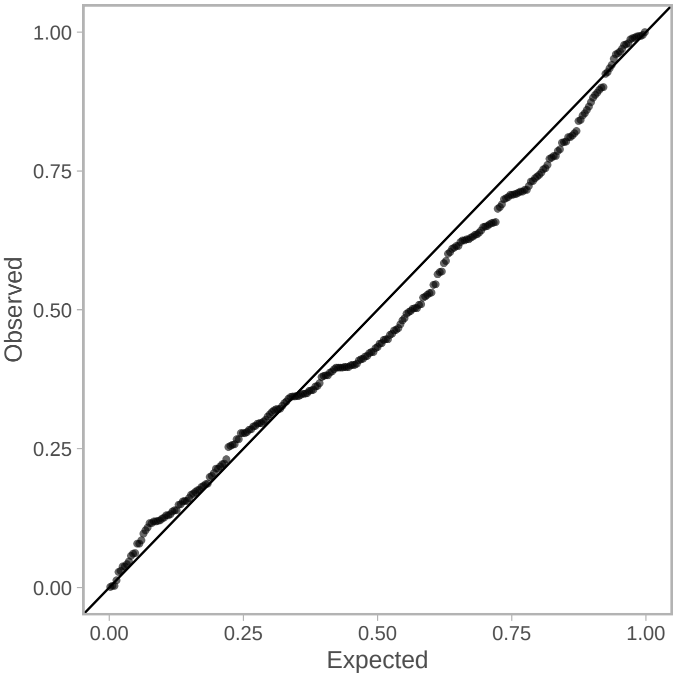


Figure S6: QQ plot of simulation-based, scaled DHARMa residuals showing the observed vs. expected quantiles under a uniform distribution from the beta model fitted to Schoener's overlap values.

### S4. Spatiotemporal relative prey weight model

Table S1: Spatiotemporal GLLM model specifications in R syntax. f refers to a factor variable, and sc that the variable is scaled.

| Model | R code |
|---|---|
| Saduria | saduria_rel_weight ~ 0 + fyear + fquarter + depth_sc + oxygen_sc + pred_length_cm_sc + small_cod_density_sc*density_saduria_sc + flounder_density_sc*density_saduria_sc |
| Benthic prey | benthic_rel_weight ~ 0 + fyear + fquarter + depth_sc + pred_length_cm_sc + small_cod_density_sc*oxygen_sc + flounder_density_sc*oxygen_sc |
| Total prey | tot_rel_weight ~ 0 + fyear + fquarter + depth_sc + pred_length_cm_sc + large_cod_density_sc*oxygen_sc + flounder_density_sc*oxygen_sc |

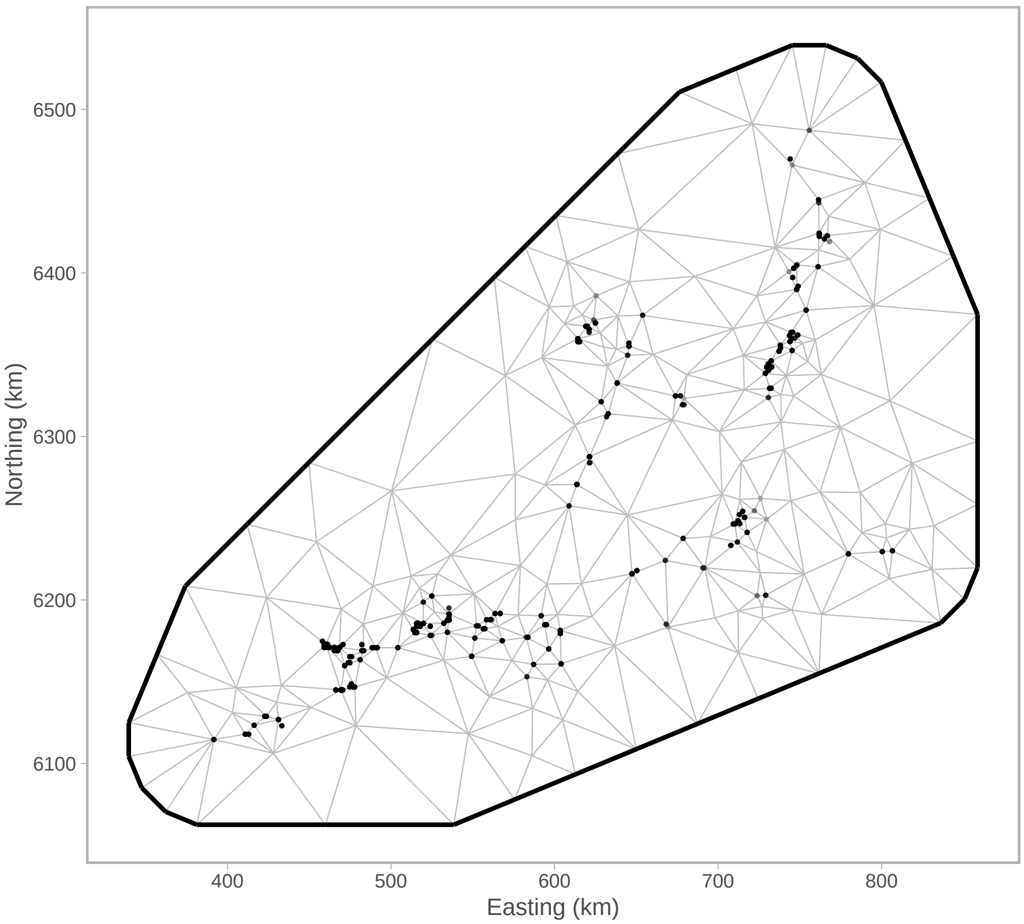


Figure S7: SPDE mesh for the flounder stomach content models.

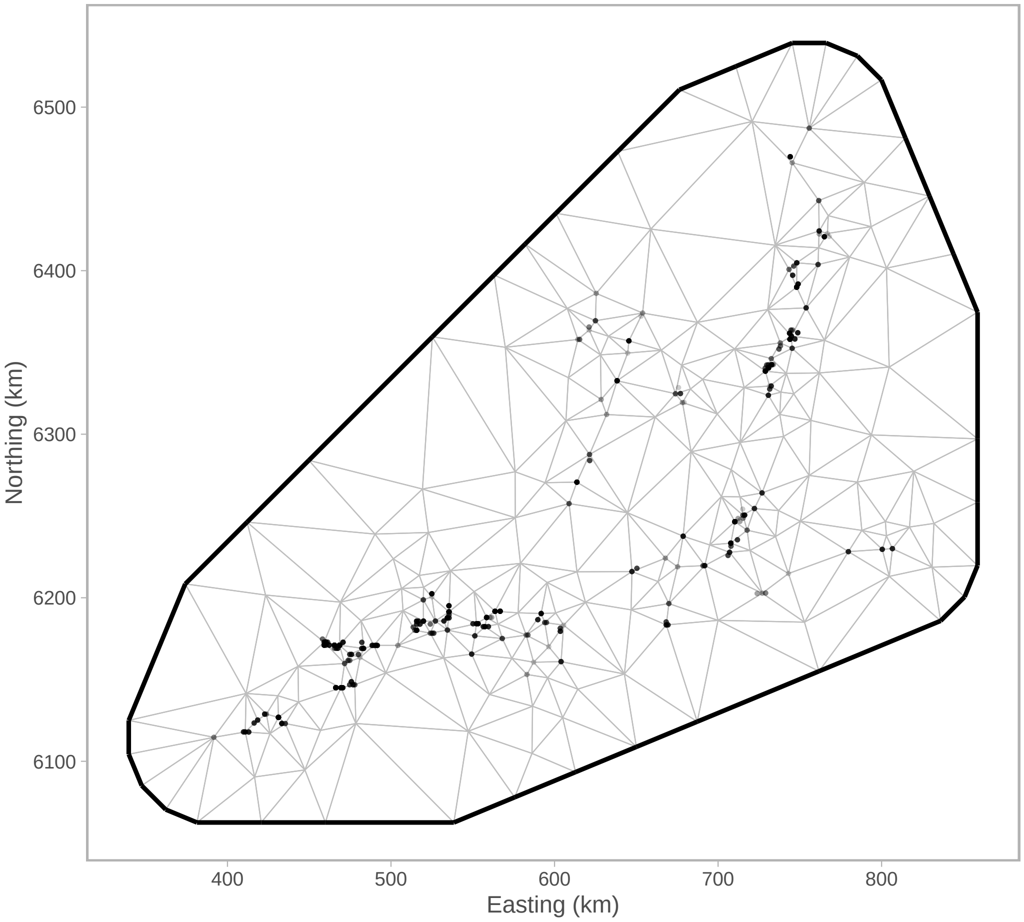


Figure S8: SPDE mesh for the small cod stomach content models.

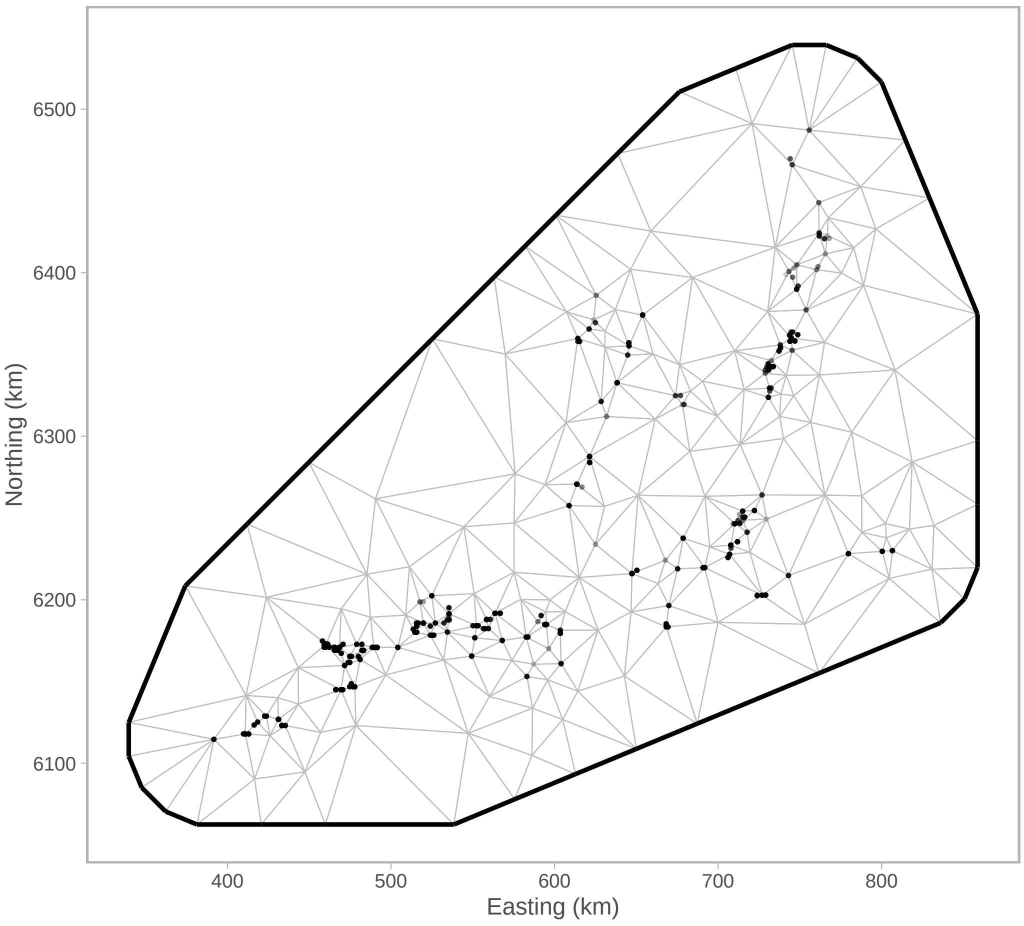


Figure S9: SPDE mesh for the large cod stomach content models.

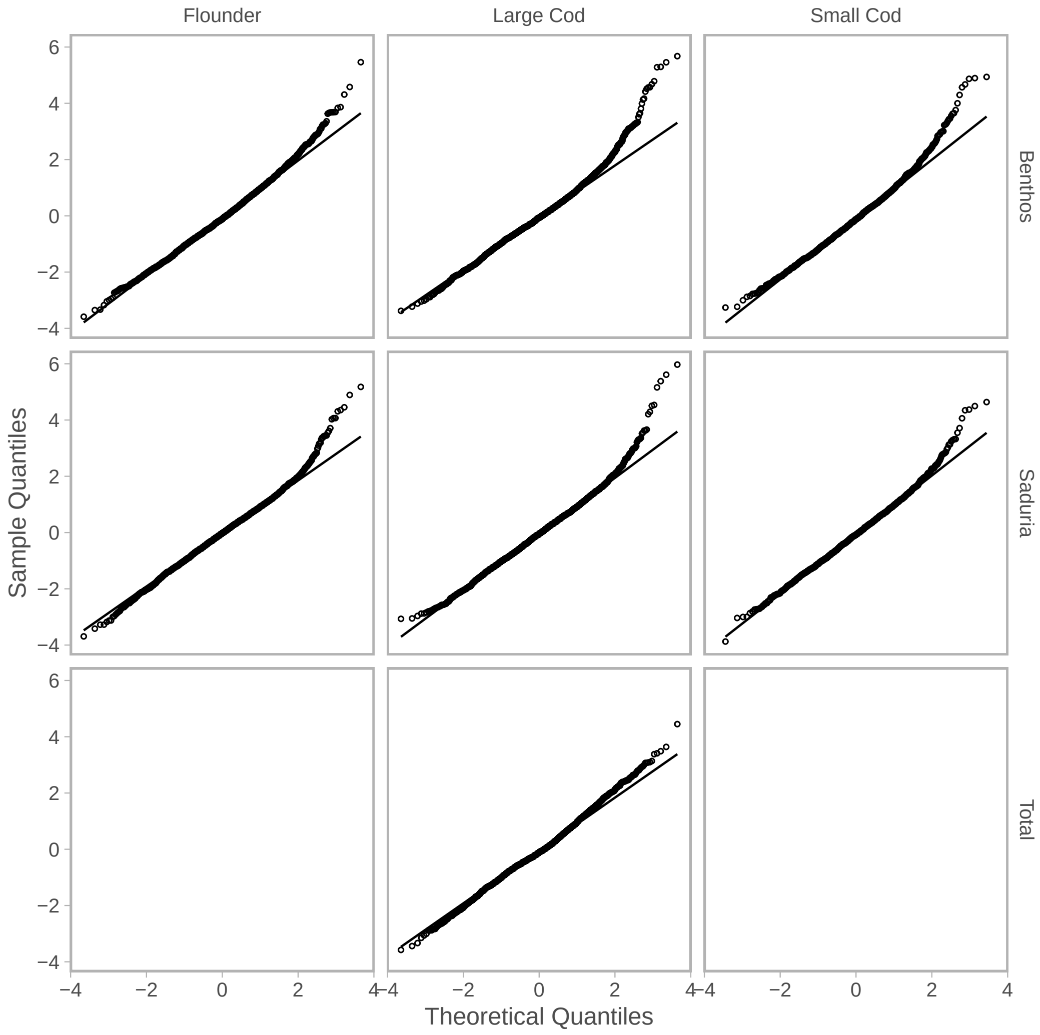


Figure S10: QQ-plots of relative prey weight models based on randomized quantile residuals, where fixed effects are held at their maximum likelihood estimate and the random effects are sampled with MCMC via tmbstan (Monnahan and Kristensen, 2018) and Stan (Stan Development Team, 2022), in line with Thygesen *et al.* (2017) and Rufener *et al.* (2021).

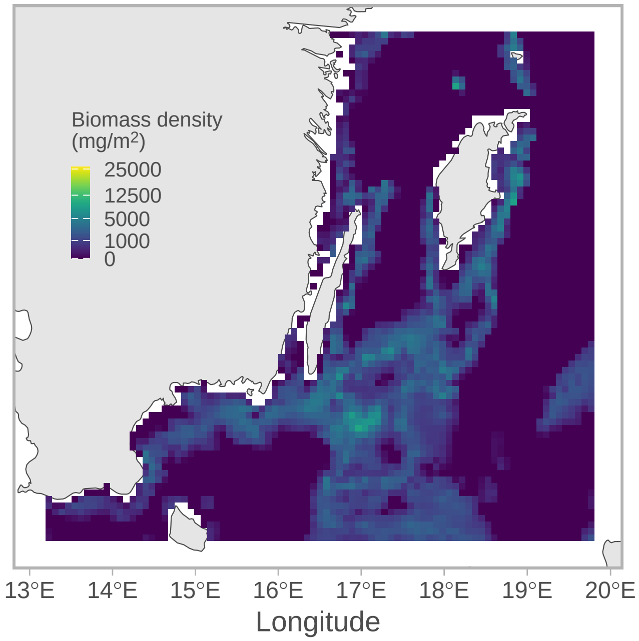


Figure S11: Predicted Saduria biomass density predictions from Gogina *et al.*, (2020).

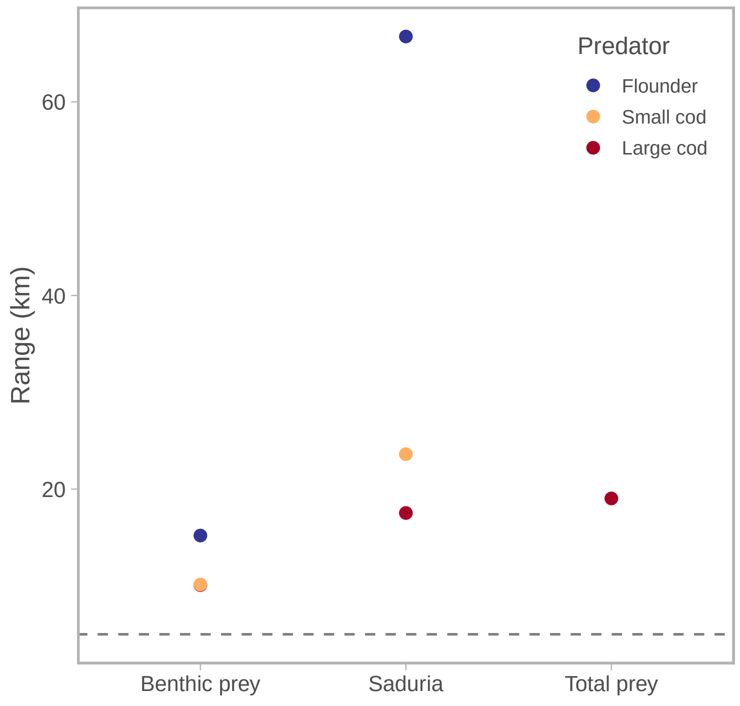


Figure S12: Spatial range estimates for spatial and spatiotemporal random fields, where color indicates the predator group. Spatial range is the distance at which correlation decays to ≈ 0.13, i.e. the distance at which two points are effectively independent. The horizontal dashed line indicates the mesh "cutoff" (minimum distance between vertices or knots).

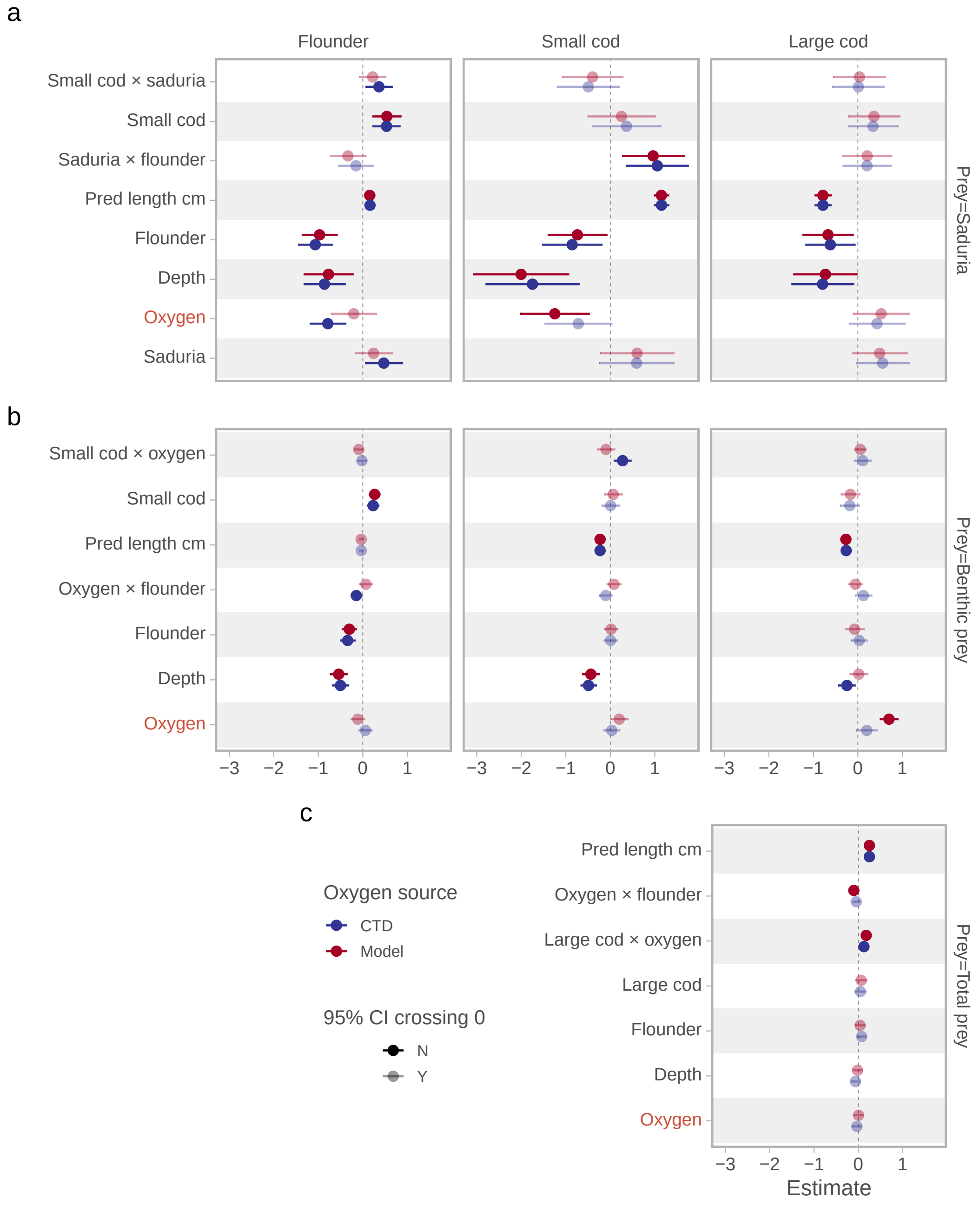


Figure S13: Sensitivity analysis of oxygen estimates (in red text) in the stomach content models. This figure depicts standardized coefficients from the spatiotemporal generalized linear mixed model fitted to relative prey weights. Predator groups are in columns, and each row represents a prey species (response variable, a=*Saduria*, b=Benthic prey, and c=Total prey weight). The y-axis shows the predictor variable. Colors indicate the oxygen source (blue=CTD, red=model based oxygen), and the transparency corresponds to the support (yes/no = Y/N) for the effect, such that semi-transparent points show estimates with 95% confidence intervals overlapping 0.

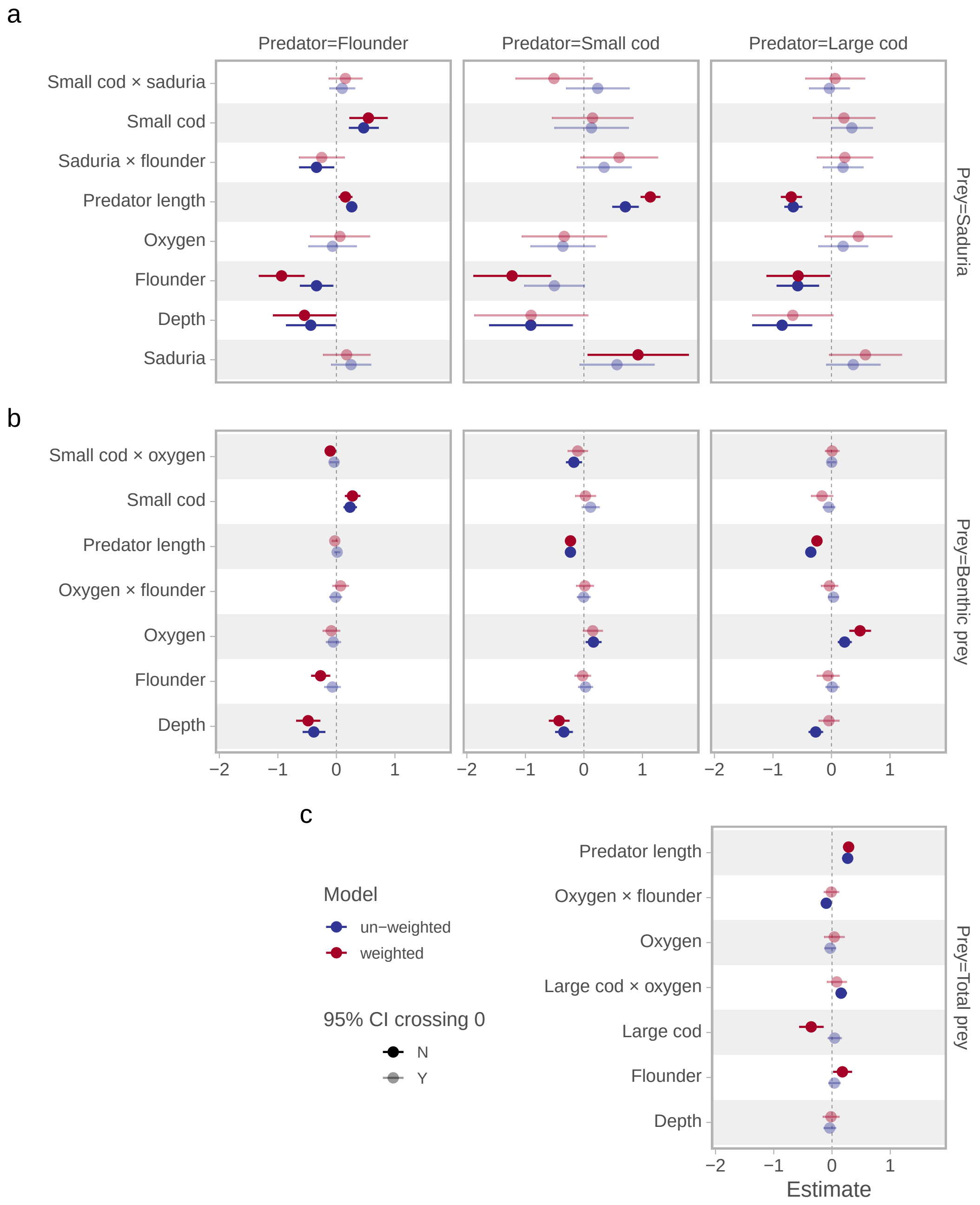


Figure S14: Sensitivity analysis of sample weights in the stomach content models. This figure depicts standardized coefficients from the spatiotemporal generalized linear mixed model fitted to relative prey weights. Predator groups are in columns, and each row represents a prey group (response variable, a=*Saduria*, b=Benthic prey, and c=Total prey weight). The y-axis shows the predictor variable. Colors indicate the model, where the un-weighted estimates are in blue and weighted estimates in red, and the transparency corresponds to the support (yes/no = Y/N) for the effect, such that semi-transparent points show estimates with 95% confidence intervals overlapping 0.

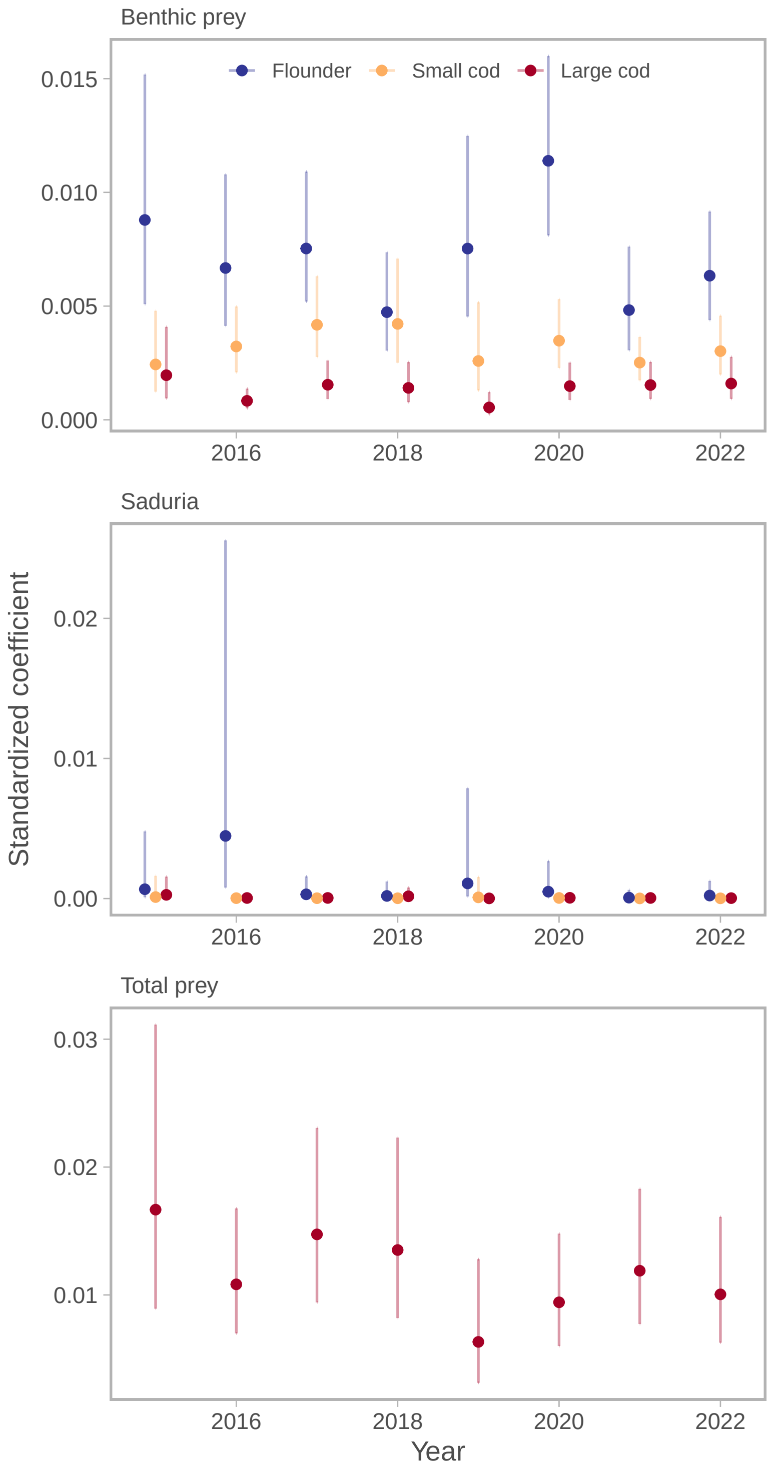


Figure S15: Effects of year on the relative prey weight of benthic prey (top), *Saduria* (middle) and total prey (bottom) in the diet. Color indicates predator group.

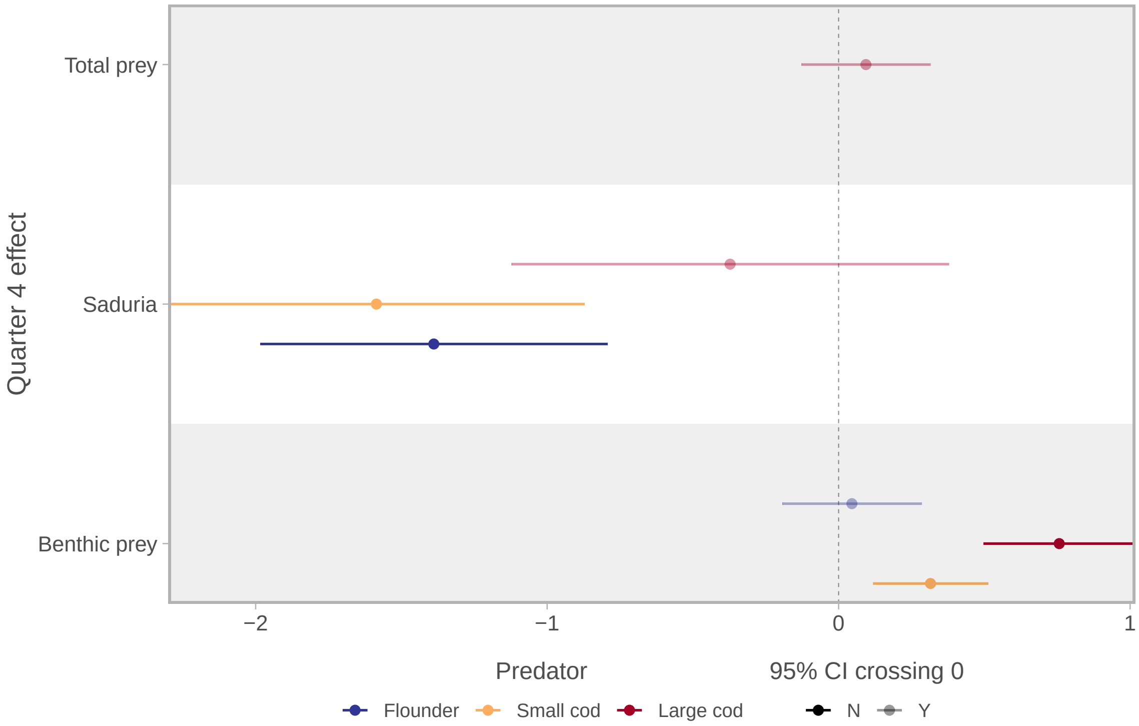


Figure S16: Effects of quarter four on the relative prey weight of benthic prey (y-axis). The estimates (x-axis) are differences between the fourth and the first quarter. Color indicates predator group.

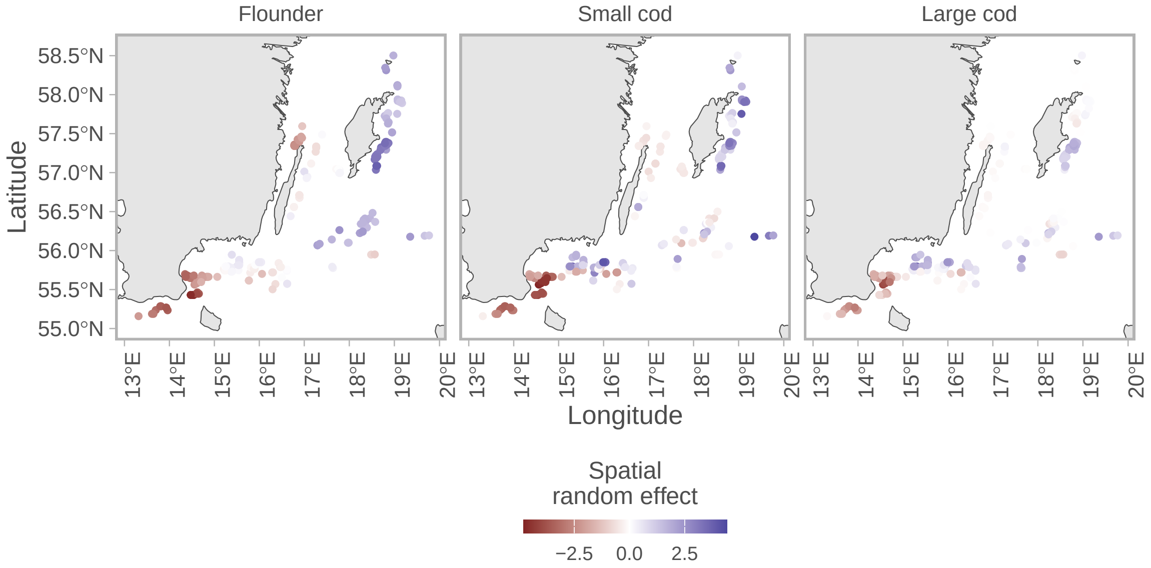


Figure S17: Spatial random effects for the *Saduria* models for all predators in link (log) space.

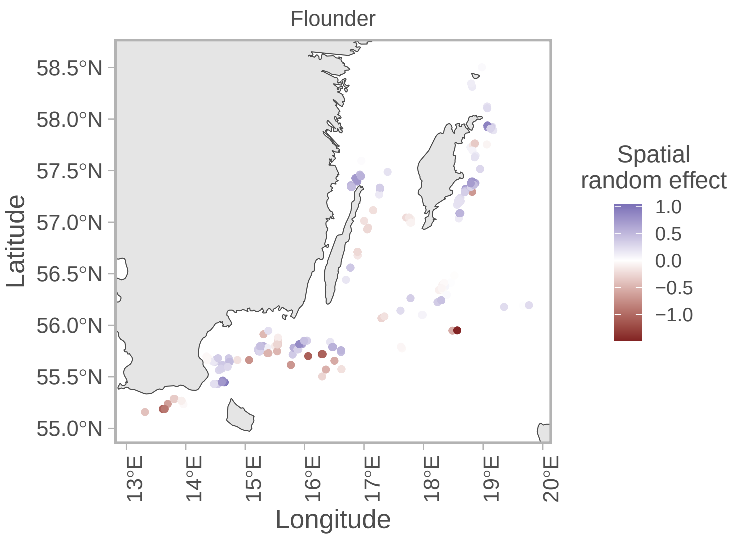


Figure S18: Spatial random effects for the benthic prey model for flounder predators in link (log) space.

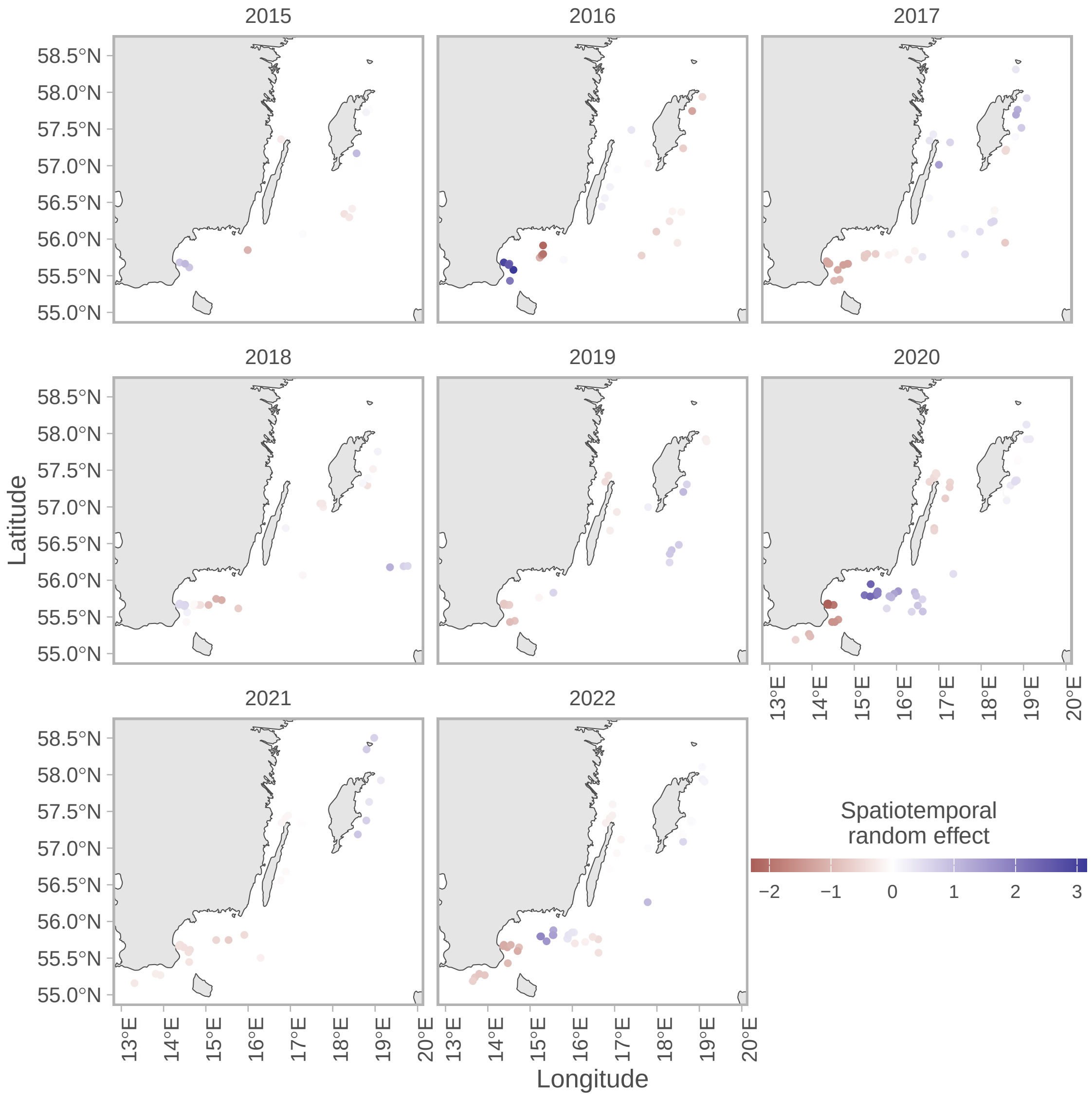


Figure S19: Spatiotemporal random effects for *Saduria* weights for flounder predators, as an example, in link (log) space.
